# The down regulation of megalin/LRP2 by transforming growth factor beta (TGF-ß1) is mediated by the SMAD2/3 signalling pathway

**DOI:** 10.1101/553974

**Authors:** Felipe Cabezas, Pamela Farfán, María-Paz Marzolo

## Abstract

Megalin/LRP2 is a receptor that plays important roles in the physiology of several organs, such as kidney, lung, intestine, and gallbladder; and also in the physiology of the nervous system. Megalin expression is reduced in diseases associated with fibrosis, including diabetic nephropathy, hepatic fibrosis and cholelithiasis, as well as in some breast and prostate cancers. One of the hallmarks of these conditions is the presence of the cytokine transforming growth factor beta (TGF-ß). Although TGF-ß has been implicated in the reduction of megalin levels, the molecular mechanism underlying this regulation is not well understood. Here, we show that treatment of two epithelial cell lines (from kidney and gallbladder) with TGF-ß1 is associated with decreased megalin mRNA and protein levels, and that these effects are reversed by inhibiting the TGF-ß1 type I receptor (TGF-ßRI). Based on *in silico* analyses, the two SMAD-binding elements (SBEs) in the megalin promoter are located at positions −57 and −605. Site-directed mutagenesis of the SBEs and chromatin immunoprecipitation (ChIP) experiments revealed that SMAD2/3 transcription factors interact with SBEs to repress the megalin promoter and that they are also required for the repressing role of TGF-ß1. In addition, high concentration of albumin reduced megalin expression and promoter activation that depend on the expression of SMAD2/3. Interestingly, the histone deacetylase inhibitor Trichostatin A (TSA), which induces megalin expression, reduced the effects of TGF-ß1on megalin mRNA levels. These data show the significance of TGF-ß and the SMAD2/3 signalling pathway in the regulation of megalin and explain the decreased megalin levels observed under conditions in which TGF-ß is upregulated, including fibrosis-associated diseases and cancer.

## Introduction

Progressive fibrosis is the final stage of several chronic diseases, including diabetes, obesity, gallstone disease, liver cirrhosis, pulmonary fibrosis, and cardiovascular and muscular diseases [1, 2]. The hallmark of this condition is increased deposition of extracellular matrix (ECM) and alteration of its composition, resulting in loss of function of the affected organs and tissues. Along with abnormal ECM deposition, fibrosis also alters the function of several cell types, including epithelial cells that undergo the epithelial-mesenchymal transition (EMT), which is characterized by loss of cell adhesion, suppression of E-cadherin expression [3, 4] and expression of genes such as vimentin and alpha-smooth muscle actin (SMA) [3, 5, 6]. The EMT is thus associated with cell migration, tumour invasion and cancer. In kidney fibrosis for example, an important population of myofibroblasts originates from renal tubular epithelial cells via EMT [7].

The pro-inflammatory cytokine TGF-β1 is a notorious inducer of fibrosis and cancer [8-10]. For example, in gallbladder cancer cells TGF-β induces EMT *in vitro* [11] and *in vivo* [12, 13]. In general, TGF-β1 increases ECM production by stimulating collagen and fibronectin synthesis, and in epithelial cells it is involved in EMT [4]. The signal transduction pathway initiated by this cytokine requires activation of the serine/threonine kinase receptors TGF-ßRI and TGF-ßRII on the cell surface, resulting in their auto phosphorylation and subsequent phosphorylation of SMAD transcription factors [14]. The SMAD2 and SMAD3 proteins mediate the canonical TGF-β1 pathway [15, 16]. Under steady-state conditions, SMAD proteins exist as homo-oligomers that reside in the cytoplasm. Ligand activation causes SMAD2/3 phosphorylation by the TGF-ßRI kinase (ALK5), leading to the formation of a complex with SMAD4. The SMAD complex can then translocate to the nucleus and regulate target genes by directly binding to the promoters with SMAD-binding elements (SBEs) or by associating with transcriptional co-activators or co-repressors [14, 17-21]. In addition to activating the canonical SMAD-dependent pathway, TGF-ß1 can also activate other signalling pathways that involve c-Jun N-terminal kinases (JNK), extracellular signal–regulated kinases (ERK) and p38 [22].

Megalin/LRP2 is a 600 kDa membrane glycoprotein that plays important physiological roles during embryonic development [23-26] as well as in adulthood [27]. The members of this family are type I membrane proteins, collectively known as lipoprotein receptor-related proteins (LRPs), which act mainly at the cell surface. Megalin is a multi-ligand endocytic receptor expressed on the apical membrane of several types of epithelial cells, including proximal tubule cells (PTCs) in the kidney [28, 29], alveolar cells in the lung [30, 31], and epithelial cells of the mammary gland [32], thyroid [29], epididymis [33], colon, prostate [34], and gallbladder [35]. In addition, megalin is also present in other cell types, including glial cells [36, 37], neurons [38] and liver macrophages [39], among others.

The majority of megalin functions are related to the internalization of its ligands. The ligands for megalin expressed at the apical membrane of PTCs are proteins and metabolites present in the glomerular filtrate [27]; for megalin expressed in the gallbladder, the ligands include molecules present in the bile such as albumin and apolipoprotein-related proteins [27, 35]. The numerous megalin ligands include apolipoproteins, complexes of vitamin A, B12 and D and their corresponding transporter proteins [40], insulin[41], leptin [42], angiotensin II (Ang II) [43], connective tissue growth factor (CTGF) [44] and albumin [28]. The intracellular activation of vitamin D to its active metabolite 1, 25(OH)_2_D_3_ requires megalin-mediated internalization of a complex comprising vitamin D-binding protein (DBP)−25(OH)D_3_ and vitamin D [45, 46]. The dysfunction of megalin has been associated with different pathological conditions, including diabetic nephropathy [47-49], gallstone disease [27] and cancer [34, 50].

Megalin expression is specifically reduced in epithelial cells in different fibrosis-associated pathologies that compromise the function of organs such as kidney, liver and gallbladder [27, 47, 49, 51-56]. However, the molecular mechanisms underlying the decreased expression of this receptor remain unclear. Diabetic rats that were administered streptozotocin exhibit increased albuminuria and type IV collagen deposition, and decreased megalin levels. These effects were reversed by injection of a soluble fragment of TGF-β1 receptor type II (sTßRIIFc) [47]. TGF-β1 is involved in the regulation of megalin receptor levels in the opossum kidney cell line (OK), derived from the renal proximal tubule. This ligand causes a reduction of megalin protein levels and reduces albumin endocytosis [52]. The inhibition of MAPK/ERK, p38, protein kinase C (PKC), protein kinase A (PKA) and Phosphoinositide 3-kinase (PI3K) showed that none of these signalling pathways were involved in lowering albumin endocytosis through TGF-β1. In contrast, the use of dominant negative forms of the transcriptional factors SMAD2 and SMAD3 reduced the effect of TGF-β1 on albumin endocytosis, suggesting that these factors could be important for the regulation of megalin levels [52]. In accordance with the profibrotic effect of albumin, high concentration of this protein significantly reduced megalin expression in PTC cells [57]. Similarly, Ang II also decreased megalin expression [58], and the *in vivo* inhibition of the angiotensin receptor (AT_1)_ with losartan protected against the reduction of megalin observed in the kidneys of rats subjected to bovine serum albumin (BSA)-induced tubule-interstitial damage and proteinuria [27]. Despite efforts to identify the pathways that lead to decreased megalin levels in fibrosis-associated diseases, little is known about the molecular factors involved in these processes.

Additionally, megalin expression on the cell surface can be modified by ectodomain shedding through a mechanism involving the activity of PKC and matrix metalloproteases [59]. On the other hand, it has also been shown that in renal fibrosis the reduction of E-cadherin is in part explained by TGF-β induced proteolysis [4]. In the same way, TGF-β1 signalling has been associated with the activation of metalloproteinases such as tumor necrosis factor-alpha converting enzyme (TACE) and the subsequent shedding process of transmembrane proteins [60, 61].

In this work, we explored two mechanisms to investigate how TGF-β1 induces the reduction of megalin levels. The first one involves proteolytic processing. For the second mechanism we tested if TGF-β1 transcriptionally regulates megalin expression through the transcription factors SMAD2/3. Our results indicate that the effects of TGF-β1 are mostly transcriptional. We showed that transcription factors SMAD2/3 decrease megalin mRNA expression by binding directly to SBEs in the human megalin promoter and are required for the repression effect of TGF-β1 and also of albumin. Our results suggest that TGF-β1 and SMAD2/3 could be critical regulators of megalin expression in various pathophysiological conditions, including chronic renal disease, cholelithiasis and pulmonary diseases.

## Materials and Methods

### Reagents

Dulbecco’s modified Eagle’s medium (DMEM), Minimum Essential Medium alpha (α-MEM), L-glutamine, penicillin/streptomycin (P/S) and trypsin were acquired from Gibco (Gibco™ Life Technologies, Inc). MEM vitamins, fatty acid-free BSA, protease inhibitors (phenylmethylsulfonyl fluoride (PMSF), antipain, leupeptin, pepstatin, orthovanadate and NaF) and Trichostatin A (TSA) were acquired from Sigma (Sigma-Aldrich). Foetal bovine serum (FBS) was acquired from HyClone (HyClone Laboratories, Inc). Lipofectamine 2000, random primers, primers and RNase were acquired from Invitrogen. DNAse I, reverse transcriptase M-MuLV Reverse™, SYBR Green master mix, P*fu* DNA Polymerase and Taq polymerase were acquired from Fermentas. Other reagents included bicinchoninic acid (BCA) assays (Pierce^®^ Thermo Fisher Scientific Inc.), luciferase assays (Promega), recombinant TGF-β1 (R&D Systems), and RNASolv reagent (Omega Bio-Tek). The following antibodies were used: anti-E-cadherin antibody (BD Bioscience), anti-ß-tubulin antibody (Millipore), phospho-SMAD2 (Ser465/467)/SMAD3 (Ser423/425), SMAD2 #3122 and SMAD3 #9523 antibodies (Cell Signalling Technology) and anti-FLAG antibody (Clontech). Horseradish peroxidase-conjugated secondary antibodies and anti-rabbit and anti-mouse antibodies were acquired from Molecular Probes. Alexa Fluor 488 Rabbit, Alexa Fluor 555 Mouse and Alexa Fluor 488 Mouse secondary antibodies were obtained from Invitrogen. The polyclonal antiserum to the recombinant human megalin cytoplasmic domain (anti-megT) has been previously described [62]. The ALK inhibitor SB-431542 (Abcam) was kindly donated by Dr. Enrique Brandan, Department of Cellular and Molecular Biology, Faculty of Biological Sciences, P. Catholic University of Chile.

### Plasmids

pCMV5, pCMV5-SMAD2-FLAG, and pCMV5-SMAD3-FLAG were provided by Dr. Seong-Jin Kim (CHA University, Korea). The 1500 pGL3-basic megalin promoter (−1500PromBasic) [63] was a gift from Dr. F. Lammert (Homburg, Germany). p800LUC, which contains the PAI-1 promoter [64], was kindly provided by Dr. Rifkin; and the pCMVβ vector was obtained from Clontech.

### Cell culture and treatment

We used the LLC-PK1 epithelial cell line derived from porcine kidneys. Cells were cultured in alfa-MEM supplemented with 7.5% FBS containing 100 U/ml P/S, 2 mM glutamine and 2.5 μg/ml Plasmocin™. These cells have been previously used in studies on proximal tubule function and megalin expression and function [27, 35]. The dog gallbladder epithelial cell line dGBECs are polarized cells that express several proteins present in gallbladder, including megalin and cubilin [35]. These cells were cultured in low-glucose DMEM supplemented with 2 mM glutamine, 1% MEM non-essential amino acids solution, 1% MEM vitamin solution, 7.5% FSB and P/S. The cells were seeded at approximately 70-80% confluence in complete media for 24 h and maintained at 37 °C in 5% CO_2_. HEK293 cells were grown and maintained in high-glucose DMEM supplemented with 10% FBS containing 100 U/ml P/S and 2.5 μg/ml plasmocin. All the cultures were maintained in humid chambers. For albumin (BSA) treatments, LLC-PK1 cells were plated at 3 x 10^4^ cells/cm^2^ for 48 h and were subsequently grown overnight in low-serum medium (alfa-MEM supplemented with 0.5% FBS containing 100 U/ml P/S, 2 mM glutamine and 2.5 mg/ml Plasmocin™) and treated with fatty acid-free BSA or vehicle (water) in serum-free medium at different time points (0.5, 1, 6 or 12 h) and concentrations (0.01, 0.1, 10, or 20 ng/ml).

For TGF-β1 treatments, LLC-PK1 cells or dGBECs were plated at 3 x 10^4^ cells/cm^2^ for 48 h and subsequently grown overnight in low-serum medium. For dGBEC we used low-glucose alfa-MEM supplemented with 2 mM glutamine, 1% MEM non-essential amino acids solution, 1% MEM vitamin solution, 0.5% FSB and P/S. Cells were treated with recombinant TGF-β1 (1, 2.5, 5, or 10 ng/ml) or vehicle (4 mM HCL + 1 mg/ml BSA) in low-serum medium for different times (6, 12, 30 or 48 h). For TSA treatment, cells were plated at 3 x 10^4^ cells/cm^2^, incubated for 48 h and subsequently treated for 6 h with 10, 50, 100 or 500 nM with TSA in the presence or absence of TGF-β1 (2.5 ng/ml). For experiments aimed at inhibiting the TGF-ß1-RI receptor kinase activity, LLC-PK1 cells were plated and treated with 10 µM SB-431542 for 20 h for qPCR and 48 h for immunoblot experiments.

### Transient plasmid transfection and siRNA silencing of SMAD2/3

LLC-PK1 or HEK293 cells were plated at 3 x 10^4^ cells/cm^2^ for 48 h and subsequently transfected with FLAG-tagged SMAD2 and FLAG-tagged SMAD3 expression vectors and −1500PromBasic or p800Luc-reporter plasmids, using Lipofectamine Reagent for 6 h, according to the manufacturer’s instructions. Finally, cells were maintained for 24-48 h in complete media before cell lysis or crosslinking. The empty pCMV5 vector was used as control. SMAD2/3 silencing in LLC-PK1 cells was performed using DsiRNA synthesized by Integrated DNA Technologies (IDT). The nucleotide target sequences for pig were 5′-GGUGAAGAAGCUAAAGAAAACAGGA-3′ and 5′-UCCUGUUUUCUUUAGCUUCUUCACCAA-3′ (siSMAD2 #1) 5′-ACAGUCAUCAUGAACUCAAAGCAAT-3′ and 5′-AUUGCUUUGAGUUCAUGAUGACUGUGA-3′ (SiSMAD2 #2), 5′-GAUAUAUCAGUGAAGAUGGAGAAAC-3′and 5′-GUUUCUCCAUCUUCACUGAUAUAUCCA-3′ (siSMAD2 #3), 5′-GGAUGCAACCUGAAGAUCUUCAACA-3′ and 5′-UGUUGAAGAUCUUCAGGUUGCAUCCUG-3′ (siSMAD3 #1), 5′-ACUGUGGUCAAGGAAGAAGACAUTT-3′ and 5′-AAAUGUCUUCUUCCUUGACCACAGUGA-3′ (siSMAD3 #2), 5′-GGUCAAGGAAGAAGACAUUUUUCTC-3′ and 5′ GAGAAAAAUGUCUUCUUCCUUGACCAC-3′ (siSMAD3 #3).

LLCPK1 cells were transfected with a mix of SMAD2 siRNAs (100 nM) and a mix of SMAD3 siRNAs (100 nM) or control siRNA and Lipofectamine 2000 for 6h. The cells were used after 48h of expression.

### RNA isolation, reverse transcription and qPCR

Total RNA was extracted using an RNA-Solv isolation system following the manufacturer’s instructions. First, genomic DNA was degraded in DNAse I for 15 min at 25 °C. The RNAs were then quantified using NanoDrop equipment. The first strand of cDNA was synthesized by reverse transcription using RevertAid™ H Minus M Mulv 200 U/μl, 1 μg of RNA in 20 μl of reaction buffer and random primers, and incubation at 42 °C for 60 min. PCR was performed to verify reverse transcription using Taq polymerase and a 0.4 μl aliquot of cDNA with specific primer pairs. The sequences of the primers used for qPCR were as follows: pig megalin 5’-CTGCTCTTGTAGACCTGGGTTC-3’ (sense) and 5’-TCGGCACAGCTACACTCATAAC-3’ (antisense);

pig PAI-1 5’-CCGTCAGCAAGAGGCAGCAAGAG-3’ (sense) and 5’-TGGTGATGGGAGGAGGCAGAGC-3’ (antisense);

pig actin 5’-CCAGATCATGTTCGAGACCTTC-3’ (sense) and 5’-TCTTCATGAGGTAGTCGGTCAG-3’ (antisense);

dog megalin 5’-CCTGCCCAAGCTACCGAGCT-3’ (sense) and 5’-CGGGCCCAAAACCAGATACTC-3 (antisense);

dog GADPH 5’-GCATCGGAGGGACCCCTCAAA-3’ (sense) and 5’-CACCCGGTTGCTGTAGCCAAA-3’ (antisense). The real-time PCR reactions were conducted using SYBR Green master mix according to the manufacturer’s instructions. A 7500 real-time PCR system (Applied Biosystems) was used to measure the relative RNA levels and the data were analysed using 7500 system SDS software.

### Immunoblot analysis

Cells were lysed in lysis buffer (phosphate-buffered saline (PBS) containing 1% Triton X-100, 2 mM PMSF, 1 mM pepstatin, 2 μM antipain, 1 μM leupeptin, and 0.3 μM aprotinin) for 30 min at 4 °C. The cells were then removed using a cell scraper, and the nuclei eliminated by centrifugation at 20,000x g for 5 min at 4 °C. Protein concentrations were measured using a BCA protein assay kit. Equal amounts of protein from each extract were subjected to reducing conditions, and megalin, E-cadherin or ß-tubulin were immunodetected as previously described [35, 62]. The immunoblots were visualized using the Pierce enhanced chemiluminescence (ECL) system, and densitometric analysis was performed using the Gel Doc 2000 Gel Documentation System and Quantity One version 4 software (BioRad, CA, USA). The β-tubulin signal was used to normalize protein levels.

### Immunofluorescence

LLC-PK1 cells and dGBECs were washed twice with Ca^2+^/Mg^2+^ PBS (137 mM NaCl, 2.7 mM KCl, 10 mM Na_2_HPO_4_, 1.8 mM KH_2_PO_4,_ 1 mM MgCl_2_ and 0.1 mM CaCl_2_), fixed with 4% paraformaldehyde in PBS for 30 min at 25 °C and then immediately permeabilized with 0.2% Triton X-100 in PBS for 10 min at 25 °C. Nonspecific binding sites were blocked with 0.2% gelatin in PBS for 5 min, and the slides were then incubated with primary antibodies (anti-megT, 1:500; anti-E-cadherin, 1:500; or anti-pSMAD2/pSMAD3, 1:100) overnight at 4 °C. The coverslips were then washed with blocking buffer and incubated with secondary antibodies (Alexa Fluor 488 Rabbit, Alexa Fluor 555 Mouse and Alexa Fluor 488 Mouse; all 1:500) for 2 h at 25 °C. The coverslips were then washed with PBS and mounted on a slide with a drop of Fluoromount-G. Finally, the slides were examined using a Leica DM 2000 fluorescence microscope.

### Determination of megalin levels by FACS

For FACS analyses of total megalin, LLC-PK1 cells were plated on 6 well plates and transfected with the plasmids pCMV5 or FLAG-tagged SMAD2 and FLAG-tagged SMAD3 using Lipofectamine. The SMAD2/3-expressing cells were washed with PBS and incubated with PBS containing 1 mM EDTA for 5 min, mechanically detached, and collected by centrifugation at 700xg for 5 min. The cells were resuspended in PBS, 1% heat-inactivated FBS with saponin 0.05% and gently mixed at 4°C for 30 min to permeabilize them. Permeabilized cells were then incubated with anti-megalin and anti-FLAG antibodies at 4°C for 1h in the presence of 0.05% saponin followed by incubation with Alexa 488 or Alexa 647 secondary antibodies (Molecular Probes) at 4°C. Cells were analyzed by FACS Calibur flow cytometer (Beckton & Dickinson). The total megalin fluorescence was determined from SMAD2/3 positives cells after subtracting the corresponding blank controls.

### Quantification of megalin levels at the cell surface and degradation assay

The levels of megalin at the cell surface were measured by cell surface biotinylation. LLC-PK1 cells were grown in 6-well plates. After treatment with TGF-β1 the cells were washed three times with ice-cold PBS and then biotinylated with sulfo-NHS-LC-biotin (Pierce) to a final concentration of 0.5 mg/mL for 1 h. After biotinylation, the cells were washed three times with ice-cold PBS, and free biotin was quenched by incubation with 50 mM NH4Cl in PBS for 10 min. Cells were lysed in lysis buffer (150 mM NaCl, 20 mM Tris, pH 8.0, 5 mM EDTA, 1% Triton-X-100, 0.2% BSA and protease inhibitors) at 4 °C. Biotinylated cell-surface proteins were then adsorbed to streptavidin agarose beads for 16 h at 4 °C. Beads were washed and then the bound proteins were analyzed by SDS-PAGE followed by immunoblotting. Megalin half-life was measured in LLC-PK1 cells, cultured on 6-well plates at 5×10^6^cells/cm^2^. Cells were treated for the indicated times with cycloheximide (100 mg/ml) with or without 2.5 ng/ml of TGF-β1. After cell lysis, samples were processed for western blot analysis. Quantifications were done in Image J, by measuring the Megalin/Tubulin ratio and normalizing to time 0.

### Quantification of carboxy-terminal fragment (CTF) levels

LLC-PK1 cells were plated on 6-well plates (CTF) in serum-free medium for 10 h. Then, the medium was replaced by serum-free medium containing 10 µM of γ-secretase inhibitor DAPT (or vehicle) for 14 h. The cells were then treated with 2.5 or 20 ng/ml TGF-β1 for 10 h in the presence of DAPT. Cells were lysed in RIPA (50 mM Tris pH 8.0, 150 mM NaCl, 150 mM EDTA pH 8.0, 1% NP40, 0.5% Deoxycholate, 0.1% SDS) and both CTF and full-length megalin were detected after SDS-PAGE by western blot analyses.

### Promoter and reporter assays

LLC-PK1 cells were co-transfected using Lipofectamine 2000 with −1500PromBasic (300 ng), mutants or the p800Luc (300 ng) luciferase reporter plus FLAG-tagged SMAD2 and SMAD3 vectors or with an empty pCMV5 vector as a control. The p800Luc luciferase reporter plasmid containing the PAI-1 promoter was used as a positive control. A fixed amount (100 ng) of internal control (pCMV-β) was also co-transfected to allow for normalization of transfection efficiency. Luciferase activity was assayed in a luminometer using a Luciferase Reporter Assay System (Promega) according to the manufacturer’s instructions. Luciferase activity was normalized to determine the transfection efficiency using β-galactosidase expression. The results are expressed as folds increase of activity compared to the control vector.

### Site-directed mutagenesis

To generate single mutations in the SBEs identified in the megalin promoter (−1500PromBasic), the following primers were used: Site 1 (MutSBE1): 5′ TCTTTCCTTCATTTTTACT**<u>ct</u>**T**<u>aa</u>**GGTTGTGCTTTCCTTTTC 3′ (sense) and 5′ GAAAAGGAAAGCACAACC**<u>tt</u>**A**<u>ag</u>**AGTAAAAATGAAGGAAAGA 3′ (antisense); Site 2 (MutSBE2): 5′ TCTAAAGGGCTTTATGCAC**<u>ct</u>**T**<u>aa</u>**GGAGGGTGGGGACTGGCG 3′ (sense) and 5′ CGCCAGTCCCCACCCTCC**<u>tt</u>**A**<u>ag</u>**GTGCATAAAGCCCTTTAGA 3′ (antisense). Mutants were generated by PCR using P*fu* DNA Polymerase according to the manufacturer’s protocol. Mutations were confirmed by DNA sequencing.

### Chromatin immunoprecipitation assays

Transfected HEK293 cells (10^6^ per chromatin immunoprecipitation (ChIP) assay) were cross-linked in suspension for 8 min at 25 °C in PBS containing 1% formaldehyde before quenching with 1.25 mM glycine. Cells were lysed in lysis buffer (50 mM Tris-HCl pH 8.0, 10 mM EDTA, 1% SDS, 1 mM PMSF, a broad-range protease inhibitor cocktail and 20 mM butyrate) for 30 min at 4 °C on a rotator, and sonicated 5 times for 30 sec on ice at 30% power (cycle 0.5) to generate chromatin fragments of ∼200–500 bp. After sedimentation, chromatin was diluted 10-fold in RIPA buffer (10 mM Tris-HCl pH 7.5, 1 mM EDTA, 0.5 mM EGTA, 1% Triton X-100, 0.1% Na-deoxycholate, and 140 mM NaCl) containing a protease inhibitor cocktail, 1 mM PMSF and 20 mM butyrate and then incubated on a rotator overnight at 4 °C with antibodies pre-coupled to beads in A/G protein agarose (50 μl A/G protein plus 10 μg FLAG antibody incubated at 4 °C for 3 h). An unrelated HA antibody was used as control. ChIP material was collected and washed 3 times in 1 ml ice-cold RIPA buffer. Crosslinking was reversed, and DNA was eluted for 6 h on a shaker at 37 °C in elution buffer (50 mM NaCl, 20 mM Tris-HCl, pH 7.5, 5 mM EDTA, and 20 mM butyrate) containing 1% SDS and 50 μg/ml proteinase K. The DNA was isolated by phenol-chloroform extraction and ethanol precipitation. The purified DNA was amplified by qPCR to detect the SBE promoter sequences.

The primers used were as follows: for the megalin SBE promoter, (−605 FW) 5’-CGCCTCTTTCCCTTTCTTTC-3’ and (−605 RV) 5’-ACGAGGAGGAAAGTCAAGAAT-3’ and (−57 FW) 5’-GATTCCCGCATGCTTGTT-3’ and (−57 RV) 5’-GAGGCATCCCGTTTTCTACC-3’; and for the PAI-1 SBEs promoter, (−580 FW) 5’-GCAGGGATGAGGGAAAGACC-3’ and (−580 RV) 5’-AACAGCCACAGGCATGCAG-3’ and (−280 FW) 5’-AGCTGAACACTAGGGGTCCT-3’ and (−280 RV) 5’-GCCTGAGAACCTCCCTTGAC-3’. These latter PCR primers were designed according to the reported sequence data for the PAI-1 promoter [65].

### Nuclear Extracts

Cells were washed twice with PBS and centrifuged for 10 min at 300x g. The pellet was resuspended in 1 ml of hypotonic lysis buffer and incubated for 30 min at 4 °C. Nuclei were centrifuged at 10,000xg for 10 min at 4 °C, and histones were extracted with 400 μl of 0.2 M sulphuric acid (H_2_SO_4_) and incubated in a rotator for 30 min at 4 °C. Eluted immunoprecipitated samples were precipitated for 30 min on ice in 25% (v/v) trichloroacetic acid (TCA) followed by centrifugation for 10 min at 16,000xg and two washes with ice-cold acetone. Samples were air dried and dissolved in sample buffer, and the protein concentration was then measured by BCA assay. The nuclear proteins were frozen in liquid nitrogen and stored in aliquots at −80 °C.

### *In silico* analysis

To locate the transcription factor binding sites in the human megalin promoter, tools from MatInspector software (Genomatix) were used as described above [66, 67].

### Statistical analysis

Immunoblots were quantified using ImageJ 1.45s software, and qPCR analysis was performed using a relative quantification mathematical model, as previously described [68]. Data are expressed as the means ± SD from at least three independent experiments. Comparisons were performed using analysis of variance (ANOVA) followed by Bonferroni correction. All of the statistical analyses were performed using GraphPad Prism 5.

## Results

### The decrease in megalin mRNA and protein levels induced by TGF-β1 in epithelial cell lines is dependent on the activation of TGF-ß type I receptor kinase

Although it has been established that TGF-ß1 reduces megalin protein levels [47, 52], it is not known whether TGF-ß1 directly regulates megalin transcription. Therefore, megalin mRNA levels were measured by qPCR in two cell lines that express megalin endogenously: LLC-PK1 cells, a cell line derived from pig kidney proximal tubules and dGBECs derived from dog gallbladder epithelial cells. The cells were treated with TGF-ß1 for different times and at different concentrations (Fig 1). TGF-ß1 concentrations of 2.5 ng/ml and 10 ng/ml significantly decreased megalin mRNA levels by 40% in LLC-PK1 cells, while treatment with 10 ng/ml TGF-ß1 decreased megalin mRNA levels by 60% in dGBECs (Figs 1A and 1C). These results indicate that both cell types exhibit a reduction in megalin mRNA expression following TGF-ß1 stimulation. At lower TGF-ß1 concentrations, only LLC-PK1 cells showed decreased megalin mRNA levels. However, megalin mRNA levels show the largest decrease (as percentage) in dGBECs when exposed to higher TGF-ß1 concentrations. Next, we studied the effects of TGF-ß1 exposure over time in LLC-PK1 cells and dGBECs by incubating them with 2.5 ng/ml and 10 ng/ml TGF-ß1 respectively. In both cell types, megalin mRNA levels significantly decreased after 6h of TGF-β1 treatment (Figs 1B and 1D). mRNA expression of plasminogen-activator inhibitor (PAI-1), a known TGF-ß1 target gene, was measured as positive control (Figs 1E and 1F) to show that the cells had an active, operative TGF-ß1 signalling pathway.

**Fig 1.**
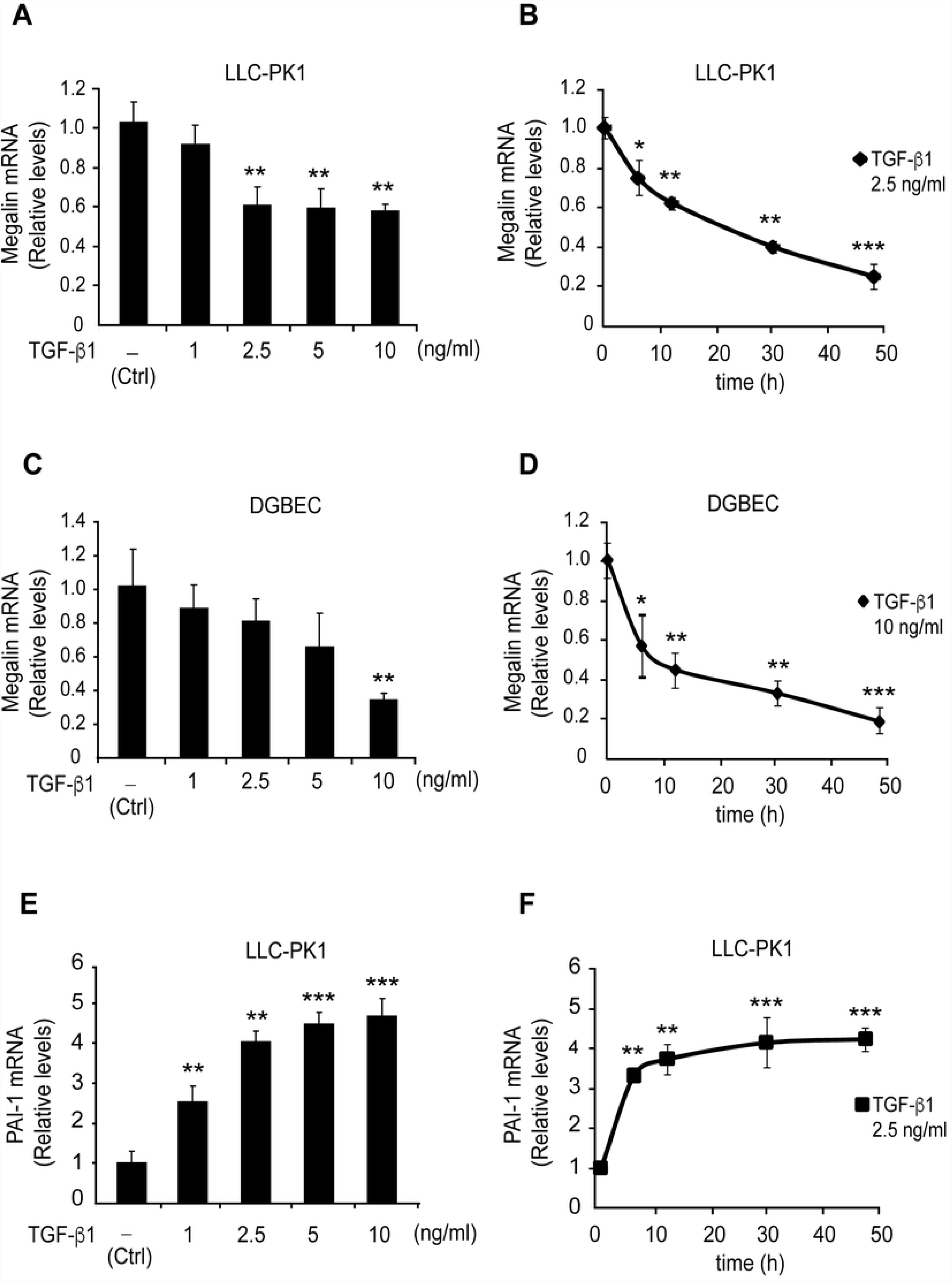
Effect of TGF-ß1 treatment on megalin mRNA levels. (A) LLC-PK1 cells and (C) dGBECs were treated with the indicated TGF-ß1 concentrations for 12 h, and megalin mRNA levels were analysed by real-time PCR. Megalin mRNA levels analysed by real-time PCR after treatment of LLC-PK1 cells (B) and (D) dGBECs with 2.5 or 10 ng/ml TGF-ß1 respectively for different times (6, 12, 30 and 48 h). As control, we measured PAI-1 mRNA levels in LLC-PK1 cells (E) that had been treated with TGF-ß1 at the indicated concentrations for 12 h or (F) treated with 2.5 ng/ml TGF-ß1 for different times (as shown in B and D). Values were normalized using either actin or GADPH as a housekeeping gene (n = 3 independent experiments). The results are expressed as the means ± standard deviation (SD). Significant differences compared with the control (vehicle) are indicated by *P≤0.05, **P≤0.01, ***P≤0.001.

Given the effect of TGF-ß1 on megalin mRNA levels, we evaluated changes in megalin protein levels by treating LLC-PK1 cells and dGBECs with 2.5 ng/ml or 10 ng/ml TGF-ß1, respectively, for 48 h. Immunoblot and immunofluorescence analyses of megalin expression in LLC-PK1 cells (Figs 2A and 2B) and dGBECs (Figs 2C and 2D) showed that megalin expression was decreased. These results are also concordant with previous studies performed in OK cells [52]. Interestingly, the distribution of megalin in LLC-PK1 and dGBECs differs in confluent cells. In dGBECs (Fig 2D) it is clearly perinuclear and vesicular, whereas in LLC-PK1 (Fig 2B) besides some vesicular localization, megalin is also found in clusters close to the cell periphery. As control we measured the expression of the epithelial marker E-cadherin (Figs 2A, and 2C), a known TGF-ß1 target that is downregulated by the cytokine [<u>69</u>]. Taken together, these results indicate that TGF-ß1 decreases megalin mRNA levels, causing a concomitant reduction of megalin protein levels.

**Fig 2.**
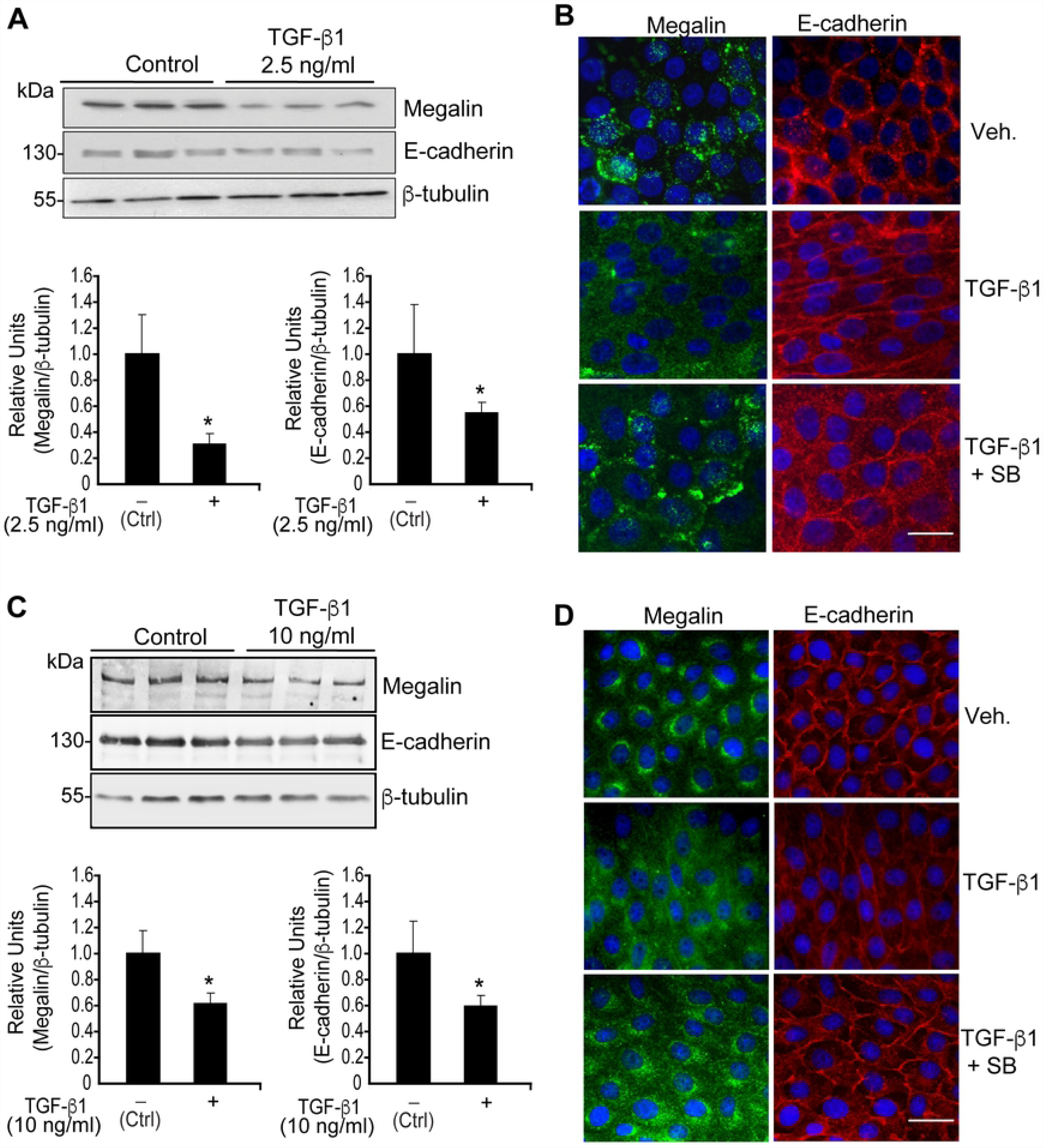
Effect of TGF-ß1 treatment on megalin protein levels. LLC-PK1 cells and dGBECs were treated with 2.5 or 10 ng/ml TGF-ß1 1, respectively, for 48 h, and total cell lysates (40 µg) from LLC-PK1 cells (A) and dGBECs (C) were subjected to SDS-PAGE and immunoblotted with antisera against megalin (top panel) or with E-cadherin antibodies (middle panel). The graphs below the immunoblots show the densitometric analyses of megalin and E-cadherin levels normalized with the housekeeping gene ß-tubulin (bottom panel). The results are expressed as the mean ± standard deviation (SD). Significant differences are indicated by *P≤0.05. Scale bars, 20 µm. (n=3 independent experiments). (B and D) Immunofluorescence for megalin and E-cadherin in confluent LLC-PK1 cells and dGBEC treated with TGF-ß1 for 48 h in the presence or absence of the inhibitor SB-431542 (10 µM). Cells were grown on coverslips, fixed in 4% paraformaldehyde and double-stained using antibodies specific for megalin (B and D; green) and E-cadherin (B and D; red). Overlapping images with Hoechst-stained nuclei (blue) are shown.

The canonical TGF-ß1 pathway includes the SMAD proteins, which are transcriptional modulators [19]. Therefore, we tested whether the SMAD2/3 signalling pathway is directly involved in the down-regulation of megalin expression. To inhibit SMAD2/3 phosphorylation (and therefore activation of the pathway), we used the specific inhibitor SB-431542 (SB), which blocks the activity of TGF-ßRI (ALK5) serine/threonine kinase. SB does not interfere with the enzymatic function of other receptors of the same family such as ALK 2,3 and 6 and does not affect the activation of other kinases (ERK, JNK or p38) associated with TGF-β pathways [70, 71]. As a result of SB treatment, both megalin and E-cadherin levels recovered from the effects of TGF-β1, as shown in the lower immunofluorescence panels (Figs 2B and 2D). When LLC-PK1 cells were treated with TGF-β1 (2.5 ng/ml) and 10 µM SB for 20 h, the effect of TGF-ß1 on megalin mRNA levels was completely reversed, and even increased compared to the control condition (Fig 3A, fourth bar). Similarly, western blot analyses (Fig 3C and 3D) show that SB completely counteracted the effect of TGF-ß1 restoring megalin protein levels. Furthermore, the addition of 10 µM SB alone resulted in a nearly 2-fold increase in basal megalin mRNA and protein levels compared with the control (Figs 3A, 3C and 3D). PAI-1 mRNA levels were measured as a response-control (Fig 3B). Interestingly, treatment with SB alone induced a significant decrease in PAI-1 mRNA levels by 30% (Fig 3B, third bar), suggesting that under steady-state conditions, LLC-PK1 epithelial cells have a basal activity of the TGF-ßRI/SMAD2/3 signal transduction pathway. In addition, to determine whether SMAD2/3 signalling responds to treatments, we used immunofluorescence to assess nuclear localization of pSMAD2/3. TGF-ß1 induced nuclear translocation of pSMAD2/3, and this effect was inhibited by SB (Fig 3E). As is shown in Fig S1, under basal conditions the cells show a diffuse SMAD2/3 staining that includes cytoplasm and nucleus. TGF-ß1 treatment for 6 h clearly defines a nuclear SMAD2/3 localization, whereas SB treatment completely abrogates the nuclear staining observed under basal control conditions, again supporting the idea that LLC-PK1 cells have a basal activity of the TGF-ßRI/SMAD2/3 pathway. Therefore, in epithelial cells, TGF-ß1 transcriptionally represses megalin expression by activating the TGF-ßRI/SMAD2/3 signal transduction pathway.

**Fig 3.**
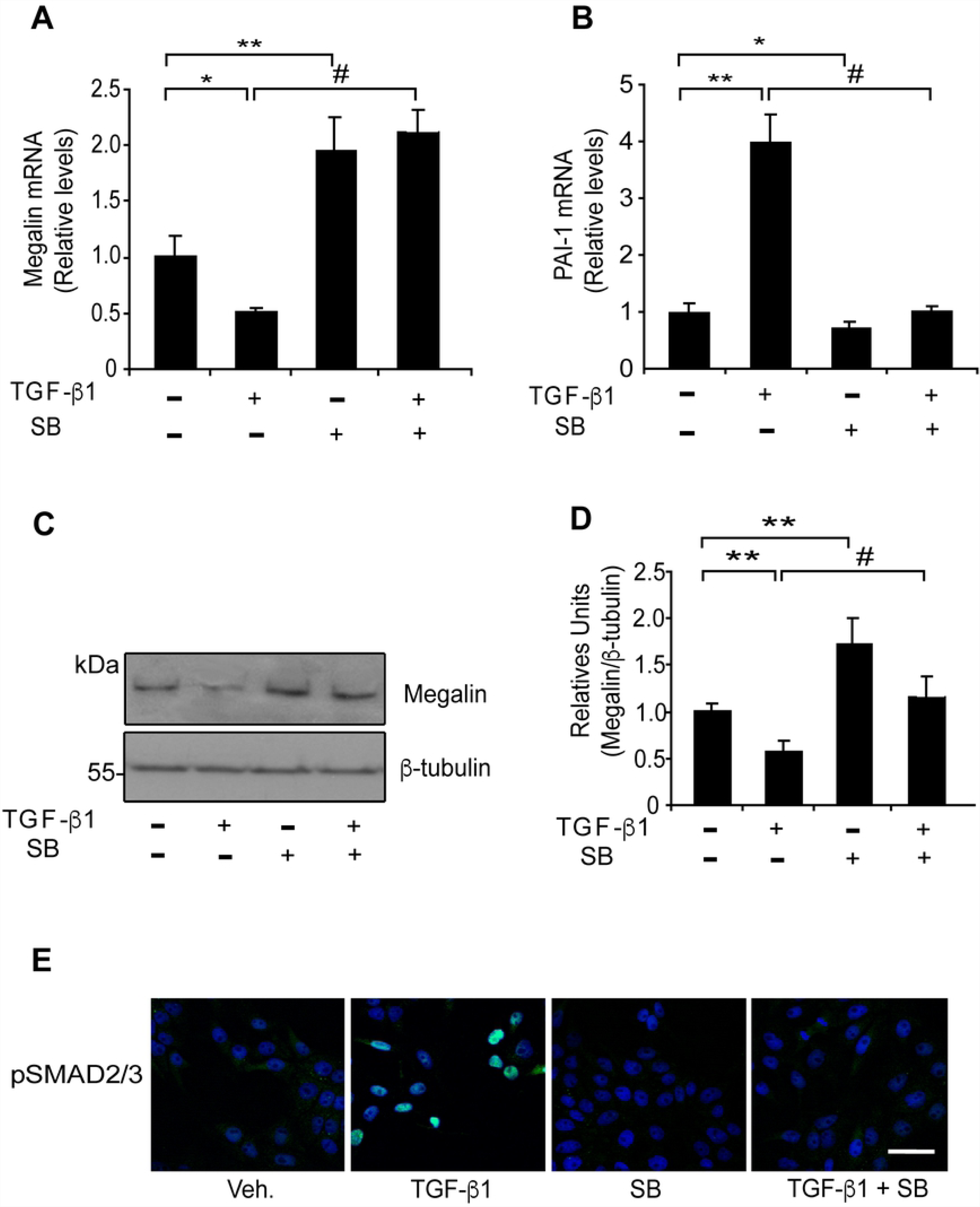
Effect of the TGF-βRI kinase activity inhibitor SB-431542 on megalin levels. LLC-PK1 cells were exposed to TGF-ß1 (2.5 ng/ml) in the presence or absence of SB-431542 (10 µM) for 20 h and megalin (A) and PAI-1 (B) mRNA levels were measured by real-time PCR. (C) LLC-PK1 cell extracts from cells treated with TGF-ß1 (2.5 ng/ml) in the presence or absence of SB-431542 (10 µM) for 48 h were used to determine megalin protein levels by immunoblotting; β-tubulin was used as the loading control. The bands in the blots were quantified by densitometry, and the results were plotted as the ratio of megalin/β-tubulin for each condition (D) Significant differences compared with the control (vehicle) are indicated by *P≤0.01, **P≤0.001. Significant differences with TGF-β1 alone are indicated by #P≤0.001 (n=4 experiments). (E) Immunofluorescence for the pSMAD2/3 transcription factors in LLC-PK1 cells. The cells were treated under each condition for 6 h (using the same concentrations described above) and stained using anti-pSMAD2/3 antibodies (green). Overlapping images with Hoechst-stained nuclei (blue) are shown. Scale bars, 20 µm.

As megalin protein levels can also be reduced by degradation, we determined if megalin half-life [72] was modified by TGF-ß1 treatment. LLC-PK1 cells were treated with TGF-ß1 2.5 ng/ml for up to 12 h in the presence of the translational inhibitor cycloheximide, in order to avoid protein synthesis and therefore follow the disappearance of megalin attributed to its degradation. Both in control and under TGF-ß1 treatment conditions, megalin protein levels decreased with very similar kinetics (Figs 4A and 4B). Previous studies showed that treating lung epithelial cells showed with a high concentration (20 ng/ml) of TGF-ß1 induces shedding of the megalin ectodomain and a reduction of megalin at the cell surface. These effects can be first detected 4h after the start of treatment and they reach a peak at 10 h [73]. Therefore, we tested if our cells respond similarly, using two different concentrations of TGF-ß1 (2.5 and 20 ng/ml) (Fig 4C). Neither cell surface megalin levels nor total levels of megalin were modified after 10 h of TGF-ß1 treatment. This result suggests that TGF-ß1 (low or high concentrations) does not induce degradation or cell surface reduction of the receptor in this cell type during this period of time (Figs 4D and 4E). Similarly, cell surface and total levels of E-cadherin were not decreased (Figs 4F and 4G).

**Fig 4.**
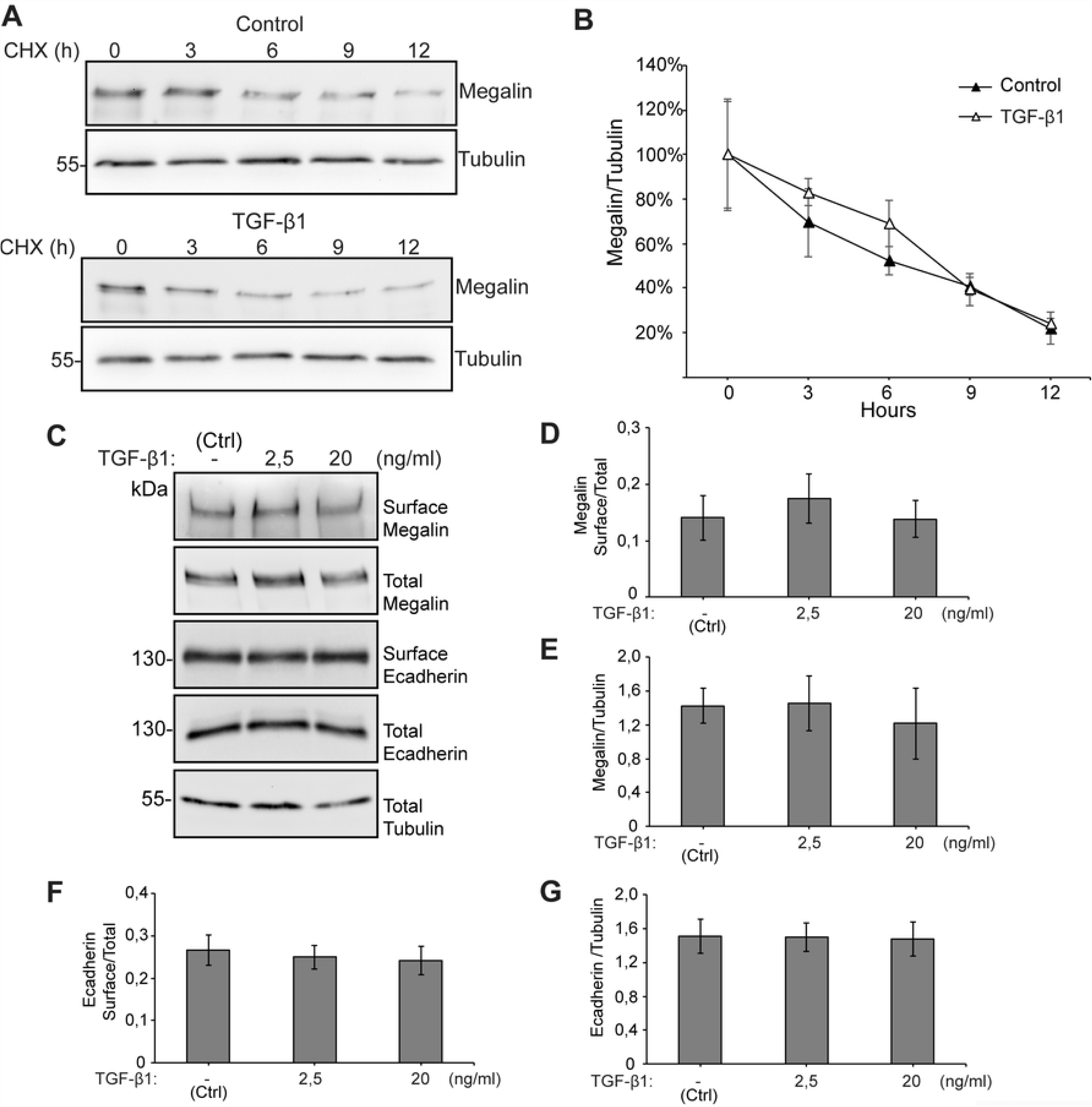
Megalin half-life and cell surface levels are not modified by TGF-ß1. (A) LLC-PK1 cells were treated with cycloheximide and either vehicle or 2.5 ng/ml TGF-ß1 for up to 12 h. At different time points, cells were lysed and the presence of megalin and β-tubulin determined by western blot. (B) Quantification of the megalin/ß-tubulin ratio (n=3 experiments). (C) LLC-PK1 cells were treated with 2.5 or 20 ng/ml TGF-ß1 for 10 h and then subjected to cell surface biotinylation at 4°C. Cell lysates were precipitated with streptavidin-Sepharose, resolved by SDS-PAGE and probed with anti-megalin or anti-E-cadherin antibodies. The graphs show the densitometric analyses of megalin and E-cadherin levels (surface/total) (D, F) as well as total protein levels (E, G). The results are expressed as the mean ± SEM (n = 4)

In order to complement this result, we also determined the levels of the megalin carboxy-terminal fragment (CTF), which is generated after ectodomain shedding. We found that CTF levels were not increased after a 10 h treatment with TGF-ß1 2.5 or 20 ng/ml. However, treatment with the γ-secretase inhibitor DAPT stabilized CTF levels (Figs 5A and 5B). The product of CTF processing is the intracellular cytoplasmic domain of megalin (MICD) which, in the case of this receptor, has been suggested to negatively control its own transcription [73]. We were able to detect the CTF but the MICD is extremely unstable and its detection erratic even in the presence of proteasome inhibitors. However, since TGF-ß1 did not increase CTF levels, we discarded an increased production of the MICD. According to the results shown in Fig 4, total megalin levels were not modified after 10 h of treatment with TGF-ß1 at the two concentrations tested (Figs 5C and 5D). Overall, these results do not support a role of TGF-ß1 in the degradation or shedding of megalin, at least in the first 10-12 h of treatment.

**Fig 5.**
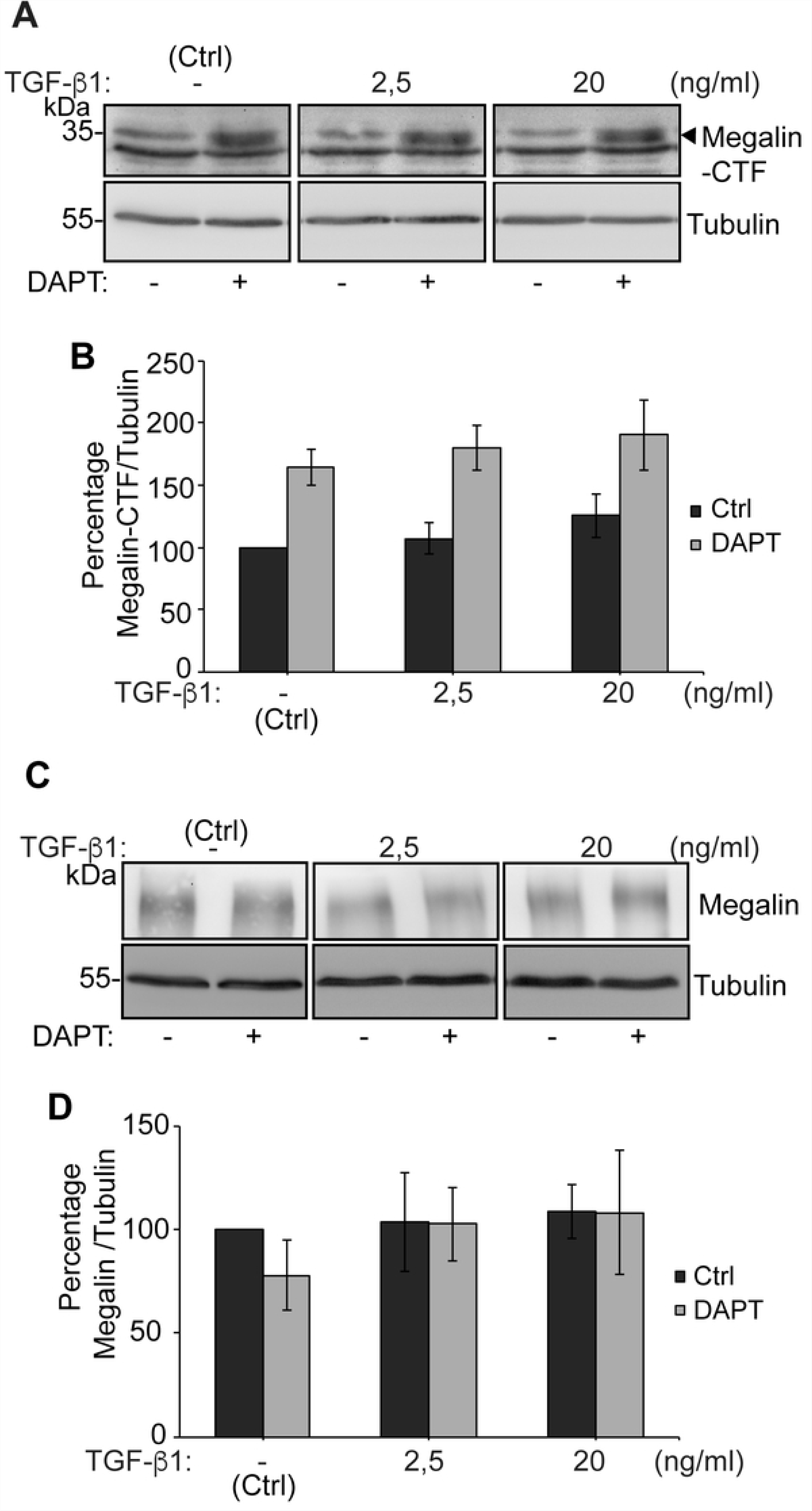
Megalin CTF levels are increased by inhibiting gamma-secretase activity but are not modified by TGF-ß1 treatment. LLC-PK1 cells were treated with DAPT 10 µM or vehicle for 24 h. During the last 10 h, cells we treated with 2.5 or 20 ng/ml TGF-ß1. (A, C) Cell lysates were processed by SDS-PAGE and probed with anti-megalin or anti-tubulin antibodies. (B, D) The graphs show the densitometric analysis of Megalin CTF/tubulin ratio and of full length Megalin/tubulin ratio with and without DAPT. The results are expressed as the mean ± SEM (n = 4).

### SMAD2/3 proteins interact with two SBEs (SMAD-binding elements) identified in the megalin promoter

The SMAD transcription factors act by forming complexes and they simultaneously activate and repress target genes in the same cell, depending on the context of the SBEs and the presence of proteins that induce the formation of the protein complexes that bind to these sites [14, 18]. We analysed the human megalin promoter with the MatInspector software and identified two consensus SBEs located at positions −605 bp and −57 of the transcription start site (Fig 6A, bold and underlined bases). In order to analyse SMAD2/3 protein-DNA interactions at the SBEs of the human megalin promoter, we performed formaldehyde crosslinking and immunoprecipitation experiments. The human megalin promoter or a parental vector was transfected into HEK293 cells in the presence of FLAG-tagged SMAD2/3, and the cells were then treated with TGF-ß1 to activate the signalling pathway and induce SMAD2/3 translocation into the nucleus. The exogenous SMAD factors were then immunoprecipitated, and the regions containing each SBE site were amplified from the complex by qPCR using two primer pairs designed for this purpose (Fig 6B). The results showed that SMAD2/3 interact with the SBEs localized at positions −605 (SBE1) and −57 (SBE2) in the human megalin promoter (Fig 6B). The relative occupancy for the factors binding to the two SBE is shown in Fig 6C (black bars). As a negative control for the immunoprecipitation experiments we used an unrelated antibody (anti-HA). The −280 and −680 SBEs from the PAI-1 promoter were analysed as positive controls for ChIP (Figs 6B and 6C, white bars). These results support the idea that SMAD2/3 could directly inhibit megalin expression at the transcriptional level by binding to the SBEs in the megalin promoter.

**Fig 6.**
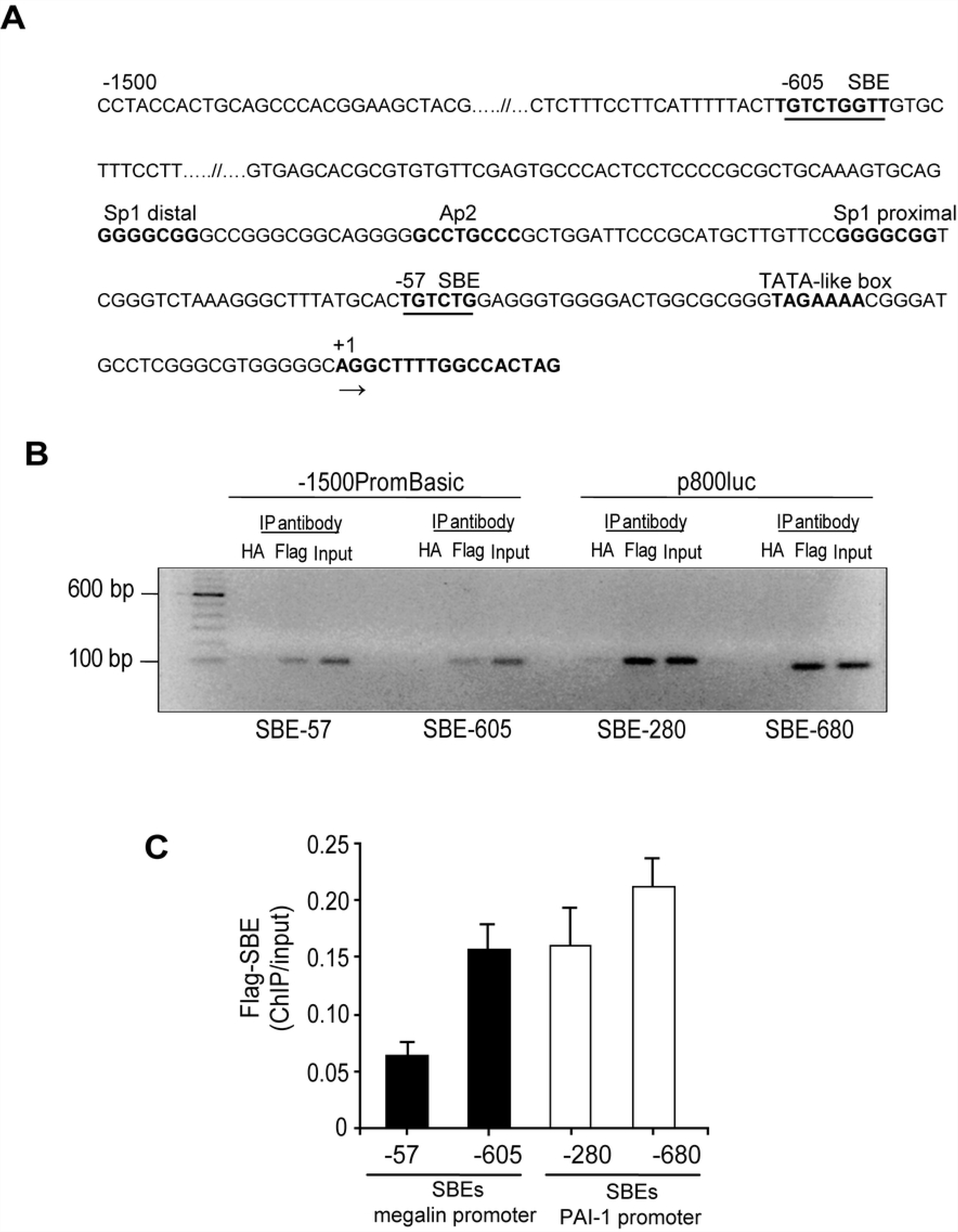
Association of SMAD2/3 with the megalin promoter. (A) Schematic representation of the human megalin promoter (−1500 bp), highlighting the putative SBEs used to analyse megalin promoter activity. (B) HEK293 cells were seeded onto 10 cm culture dishes and transfected with the megalin promoter construct (- 1500PromBasic) or the PAI-1 promoter construct (p800Luc) 48 h later. Co-transfections were performed in the presence or absence of exogenous FLAG-SMAD2 and FLAG-SMAD3 plasmids. After 24 h, the transfected cells were treated with 5 ng/ml TGF-ß1 for 1 hour and subjected to ChIP as described in the experimental section. The immunoprecipitated DNA was queried using negative control primers, a non-specific primer set and specific primers to the SBE-57 or SBE-605 sites of the megalin promoter and the SBE-280 or SBE-680 sites of the PAI-1 promoter (positive control). PCR was performed using DNA that was immunoprecipitated using an HA antibody as a negative control (HA lane). The PCR from the input DNA served as another positive control (input lane). The products were analysed by agarose gel electrophoresis. (C) The qPCR signals derived from the ChIP samples were divided by the qPCR signals derived from the input sample collected earlier during the ChIP procedure. The relative fold change was calculated from the mean ± SD 2(^)ΔΔCT values (n = 2), as described in the experimental section, and normalized with the input.

### TGF-ß1 represses megalin promoter activity via SMAD2/3

To directly test the repressive effect of SMAD2/3 on the human megalin promoter, LLC-PK1 cells were co-transfected with the human megalin promoter coupled to luciferase (- 1500PromBasic) in the presence or absence of overexpressed SMAD2/3. The results showed that the expression of SMAD2/3 triggered a significant decrease (50%) in megalin promoter activity compared with the control without SMAD2/3 transfection (Fig 7A). The positive control was the p800Luc plasmid, containing the PAI-1 promoter fragment (−800 to +71) with two consensus SBEs located at positions −280 and −680 bp of the transcription start site [64, 65]. In this case, the overexpression of SMAD2/3 increased the activation of the PAI-1 promoter (Fig 7B). Interestingly, in a complementary experiment we found that megalin protein levels were significantly decreased by the overexpression of SMAD2/3. Immunoblots showed that megalin levels decreased almost 40% in cells overexpressing SMADs (Figs 7E and 7F), and we obtained similar results when megalin levels were detected by FACS. The cells expressing SMAD2/3 with the FLAG epitope expressed significantly lower levels of megalin than the controls (Fig 7G). On the other hand, the direct effect of TGF-ß1 was also tested with the same promoters coupled to luciferase and, as expected, the cytokine decreased luciferase expression when the luciferase gene was under the control of the megalin promoter (Fig 7C). Also, as expected, TGF-ß1 increased luciferase activity in cells where the luciferase gene was under the control of p800Luc (Fig 7D).

**Fig 7.**
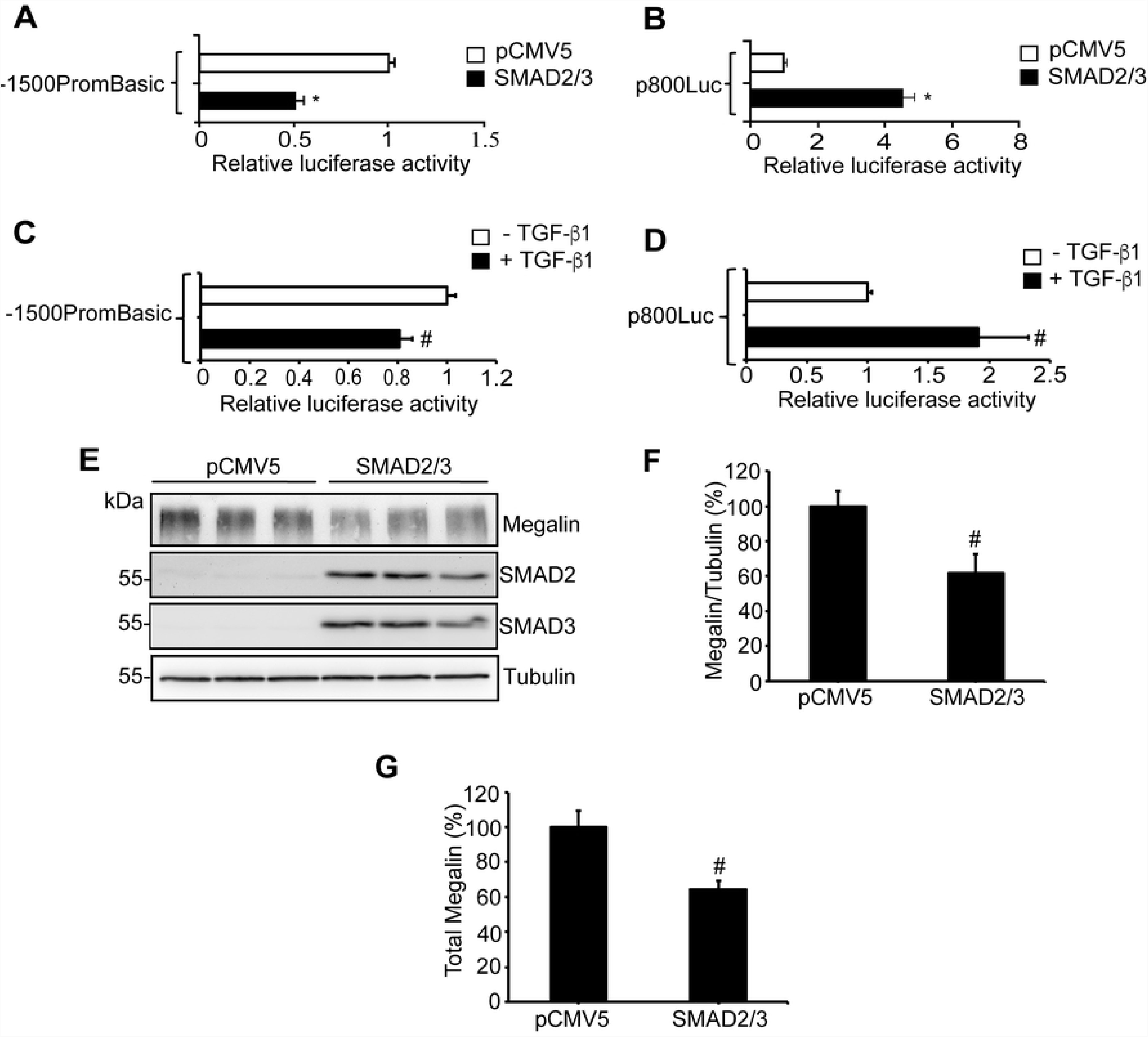
SMAD2/3 and TGF-ß1 suppress expression from the megalin promoter. (A) LLC-PK1 cells were seeded in triplicate in 24-well plates. The cells were transfected 48 h later with the megalin promoter construct (−1500PromBasic) and a ß-galactosidase vector (pCMVß). Co-transfections were performed in the presence or absence of plasmids encoding for SMAD2 and SMAD3. After 24 h, the transfected cells were lysed, and luciferase activity was measured. Cells expressing the megalin promoter construct showed a 50% decreased in luciferase activity when the SMAD transcription factors were overexpressed. (B) The overexpression of SMAD factors increased luciferase expression from the p800luc plasmid (positive control). Data in (A) and (B) are expressed as the means ± SD, and significant differences with the control (pCMV5) are indicated by *P≤ 0.001. (C) LLC-PK1 cells were transfected with the megalin promoter construct (1500PromBasic) and a β-galactosidase vector (pCMVβ). After 24h the cells were incubated in the presence or absence of TGF-ß1 2.5 ng/ml for another 24h and luciferase activity was measured in the cell lysates. Cells expressing the megalin promoter construct in the presence of TGF-ß1 show decreased luciferase activity. (D) Positive control with the p800luc plasmid in which TGF-ß1 increases luciferase expression. Data in (C) and (D) are expressed as the means ± SD, and significant differences with the control are indicated by #P≤0.05. (E) LLC-PK1 cells were transfected with pCMV5 or with SMAD2 and SMAD3 cDNAs. After 48 h of expression, the cells were lysed and megalin, SMAD2 and SMAD3 were analyzed by immunoblot; tubulin was used as a loading control. (F) Quantification of megalin/tubulin ratio in cells transfected with SMAD2 and SMAD3. (G) LLC-PK1 cells, expressing pCMV5 or SMAD2 and SMAD3 cDNAs were permeabilized with 0.1% saponin and incubated with anti-megalin and anti-FLAG. Cells were then incubated with anti-mouse Alexa−647 and anti-rabbit Alexa −488 antibodies and analyzed by flow cytometry. The results are plotted as the % of the total megalin observed in SMAD2/3 positives cells. Data in (F) and (G) are expressed as the means ± SE, and significant differences with the control are indicated by #P≤0.05.

To directly address the role of SMAD2/3 in the repressive effect of TGF-ß1 on megalin, we silenced the expression of SMAD2/3 in LLC-PK1 cells and then we transfected them with the different promoter constructs coupled to luciferase. The silencing was very efficient (84% decrease of SMAD2 and 72% of SMAD3), as seen in the immunoblot for SMAD 2 and SMAD3 (Fig 8A and 8B). Under conditions in which the expression of SMAD2/3 was significantly downregulated, the repressive role of TGF-ß1 on the activation of the megalin promoter was not evident, indicating that the cytokine affects megalin expression through the canonical SMAD2/3 pathway (Fig 8C). As control, the TGF-ß1 induced activation of p800Luc was blocked in the SMAD2/3 knock-down cells (Fig 8D). At the protein level, the downregulation of megalin induced by the cytokine was also dependent on the expression of SMAD2/3 (Fig 8E and 8F). The reduction of SMAD2/3 expression was associated with a significant decrease in the activation of the p800Luc promoter under non-stimulation conditions (Fig 8D), indicating a basal activation of the system in this cell type, similarly to what was found when TGF-βRI was inhibited by SB (Fig 3B).

**Fig 8.**
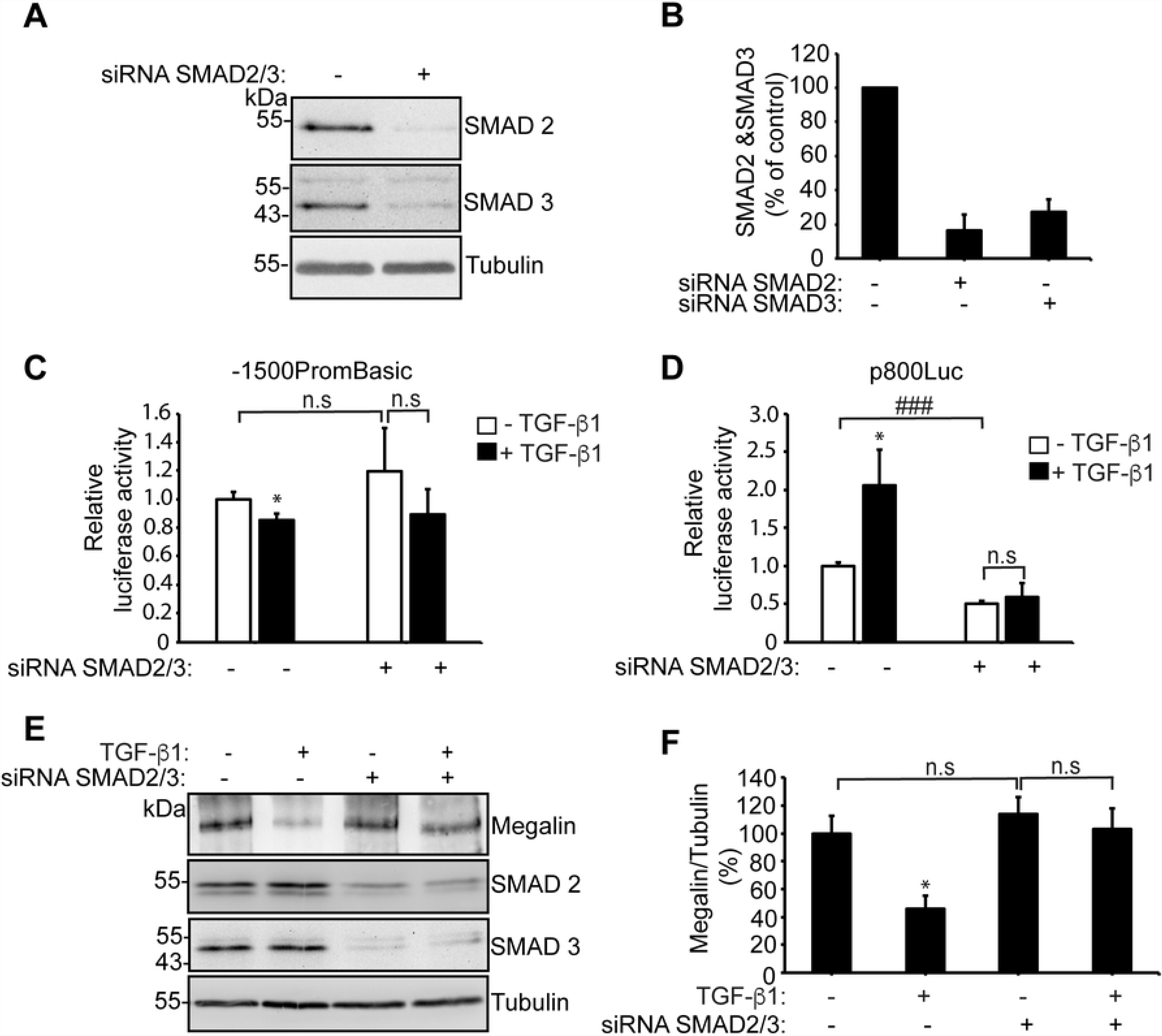
The effects of TGF-ß1 on megalin expression depend on the expression of SMAD2/3. (A) LLC-PK1 cells were transfected with control siRNA o SMAD2/SMAD3-specific siRNAs. After 48 h of expression, SMAD2 and SMAD3 were analyzed by immunoblot and tubulin was measured as a loading control (B) Quantification of SMAD2/tubulin or SMAD3/tubulin in cells transfected with siRNA. (C, D) SMAD2/SMAD3 were silenced with SMAD2/SMAD3-siRNAs in LLC-PK1 cells. The cells were then transfected with the megalin promoter construct 1500PromBasic (C) or the p800luc plasmid (D). After 24h the cells were incubated with vehicle or 2,5 ng/ml of TGF-ß1 1 for another 24h. Luciferase activity was determined using a Luciferase Reporter Assay System, and the ß-galactosidase signal was used to normalize the results. Data are expressed as the means ± SD, and significant differences with the control are indicated by *P≤0.05, while significant differences between non silenced and silenced cells are indicated by ### P≤0,001. (E, F) LLC-PK1 cells were transfected with control siRNA o SMAD2/SMAD3-siRNAs. After 24 h of expression, cells were cultured in low serum medium (0.5% FBS) for 16 h and then treated in the presence or absence of 2.5 ng/ml of TGF-ß1 or 24h. After 60h of silencing and 24h of treatment with TGF-ß1, the cells were lysed and the analyzed by immunoblot. In (E) the expression of megalin, SMAD2 and SMAD3 were determined and the detection of tubulin was used as a loading control. (F) Shows the quantification of megalin/tubulin, as % of the control, in cells transfected with siRNA and treated with TGF-ß1. Data are expressed as the means ± SE. Significant differences with the control are indicated by *P≤0.05.

Taking into account the presence of two SBEs in the megalin promoter, we established the contribution of each one in the repression of the promoter. SBE1 and SBE2 were mutated by base substitution (Fig 9A, asterisks). The plasmids were then transfected into LLC-PK1 cells in the presence or absence of overexpressed SMAD2/3, and their activities were measured. The single-site SBE mutants (Mut-SBE1 or Mut-SBE2) were able to respond to the presence of SMAD2/3 decreasing the expression of luciferase (Fig 9B, black bars compared with white bars). However, the double-mutant SBEs (MutSBE1/2) did not respond to the repressive effect of SMAD2/3 (Fig 9B, black bars compared with white bars). These data indicate that the two SMAD response elements independently repress the megalin promoter. In addition, in the SBE mutant promoters, the basal activity was significantly increased compared with the wild type promoter in the absence of overexpressed SMAD factors (Fig 9B, all white bars), suggesting that endogenous SMAD factors were basally active in LLC-PK1 cells (as suggested in Fig 3A, 3C, 3D and FigS1). Finally, the relevance of these two SBEs in the repressor effect of TGF-ß1 was determined by the same type of assays using the wild-type and mutant promoters (Fig 9C). In the SBE1 mutant and in the double SBE1/2 mutant, the repressor role of TGF-ß1 was abolished. The mutant SBE2 responded to the cytokine but to a lesser extent than the wild-type promoter. Overall, these results strongly suggest that both SBEs identified in the human megalin promoter could function as binding sites for a transcriptional repressor complex that is linked by SMAD2/3 and have a role in the repressive effect of TGF-ß1.

**Fig 9.**
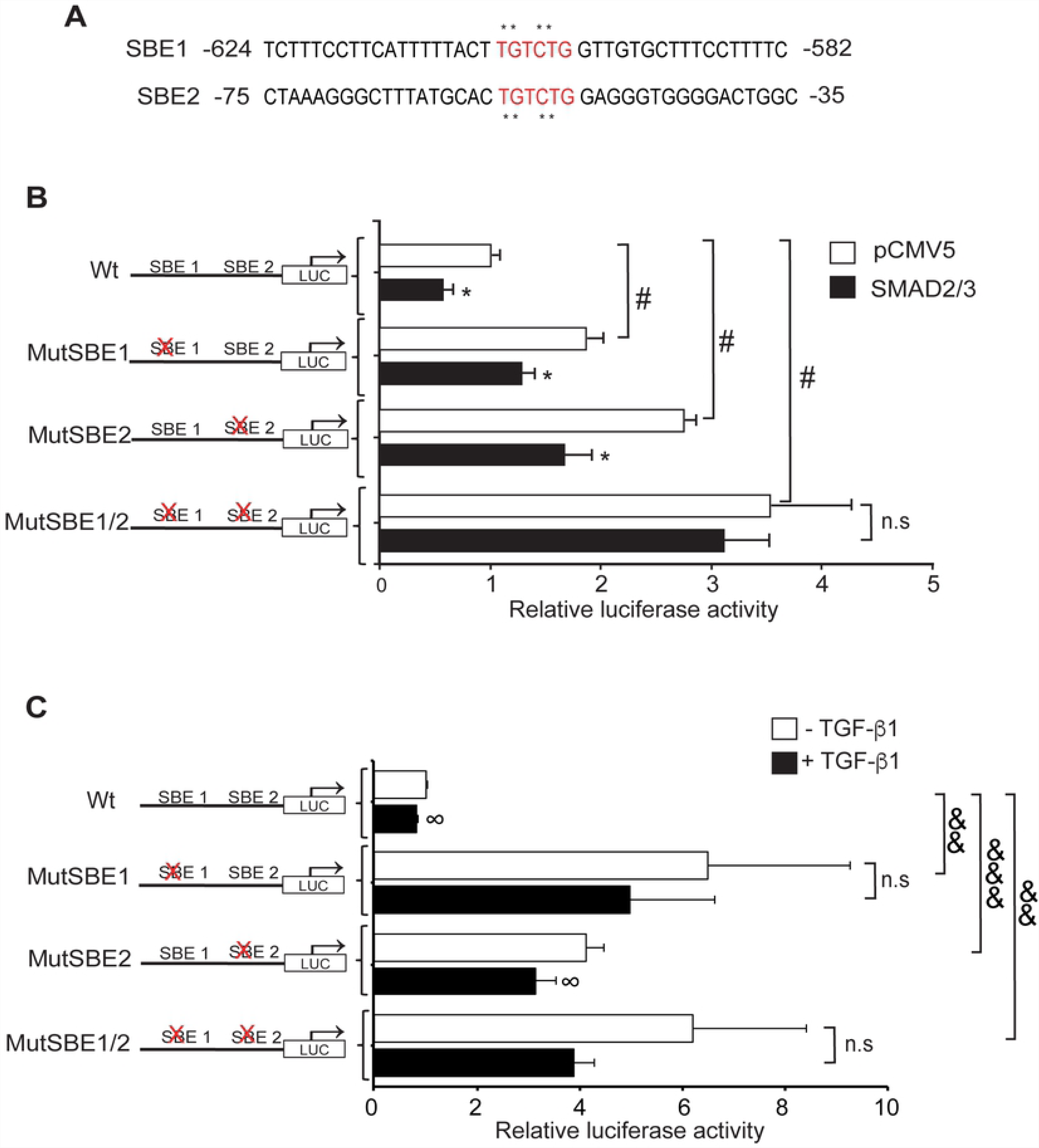
SMAD2/3 suppress megalin promoter activation at SBEs. (A) Schematic representation of SBEs and site-directed mutations (asterisks) generated in the human megalin promoter constructs. (B) LLC-PK1 cells were transfected with wild type or mutant SBE constructs of the human megalin promoter (1500PromBasic) and with the β-galactosidase vector (pCMVβ). Co-transfections were performed in the presence (black bars) or absence (white bars) of SMAD2 and SMAD3 cDNAs. After 24 h, the transfected cells were lysed, and luciferase activity was measured. Luciferase activity was normalized with ß-galactosidase expression to determine transfection efficiency. The results are expressed as the means ± SD. Significant differences with the respective controls (untransfected SMAD factors) are indicated by *P≤ 0.001. Significant differences with the wild type construct are indicated by #P≤0.001. (C) LLC-PK1 cells expressing wild type or mutant SBE constructs of the human megalin promoter (1500PromBasic) and β-galactosidase vector (pCMVβ) were incubated in the presence or absence of 2.5 ng/ml of TGF-ß1 for 24h. Later, the cells were lysed, and luciferase activity was measured. The results are expressed as the means ± SD. Significant differences with the respective controls are indicated by ∞P≤ 0.05. Significant differences with the wild type construct are indicated by &&P≤0.01, &&&P≤0.001.

### Albumin reduces megalin expression by a mechanism that involves the TGF-βRI/SMAD2/3 pathway

Although high concentrations of albumin are known to decrease megalin expression [57], it is not known if this effect is dependent on the activation of the TGF-β signalling pathway. Albumin induces TGF-ß1 secretion [74] in human proximal tubule HK-2 cells and in cells from the collecting duct that lack megalin [75]. Therefore, the pro-inflammatory effects of albumin could be megalin independent. To determine whether the albumin-induced megalin reduction is connected to the canonical TGF-ß1 pathway, LLC-PK1 cells were incubated with increasing concentrations of albumin for 24 h and the mRNA expression of megalin and PAI-I was measured. As previously reported, albumin significantly reduced megalin expression at a concentration of 10 mg/ml [57] (Fig 10A). A lower concentration was able to induce PAI-1 expression (Fig 10B). Moreover, the albumin-induced decrease in megalin expression and the increase in PAI-1 expression were significantly overcome by SB (P ≤ 0.05 in both cases), suggesting that the effects of albumin involves, directly or indirectly, activation of the TGF-ß1 receptor (Figs 10C and 10D). Again, we observed that the inhibition of the TGF-ßRI by SB significantly increased megalin mRNA levels and reduced PAI-1 expression as shown before (Fig 3A and B). To complement these results, the effects of albumin were determined directly in the activation of megalin human promoter as described for TGF-β1 (Fig 7 and Fig 8). Similarly to the effect of the cytokine, albumin significantly decreased megalin expression (Fig 10E). Moreover, the repressing effect of albumin on megalin was dependent on the presence of SMAD2/3 (Figure 10F) since it was abolished in SMAD2/3 knockdown cells.

**Fig 10.**
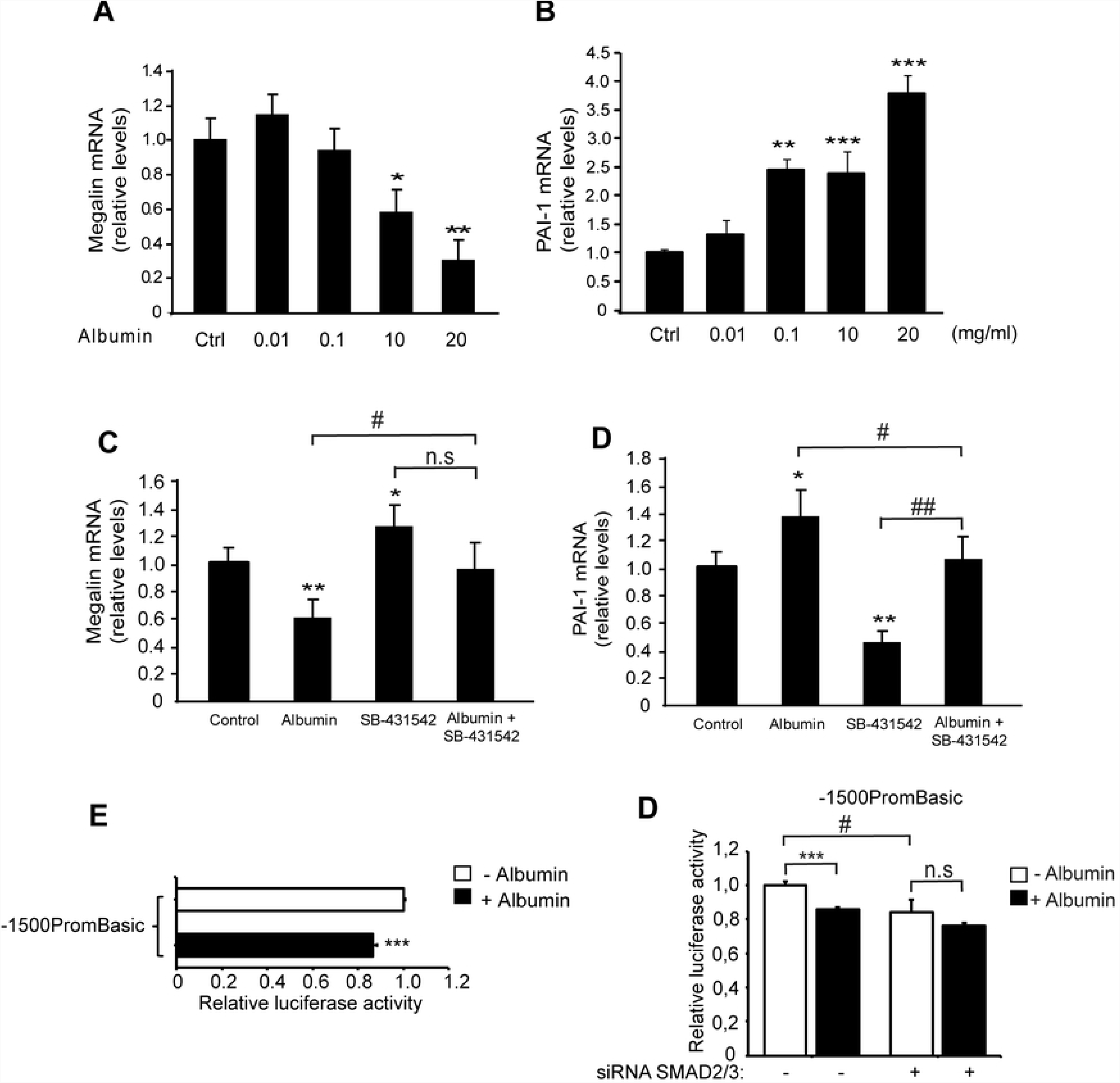
Albumin decreases megalin expression and this effect is counteracted by TGF-βRI inhibition and SMAD2/3 silencing. LLC-PK1 cells were treated with albumin concentrations ranging from 0.01 to 20 mg/ml for 24 h. Megalin (A) and PAI-1 mRNA levels (B) were measured by qPCR. (C) Megalin and (D) PAI-1 mRNA levels in LLC-PK1 cells that were exposed to 20 mg/ml of albumin and/or the inhibitor SB for 20 h. The results are expressed as the average RQ ± standard deviation (SD). (E) LLC-PK1 cells were transfected with the megalin promoter construct (1500PromBasic) and a β-galactosidase vector (pCMVβ) and incubated in the presence or absence of 20 mg/ml of albumin for 24h. Later, the cells were lysed, and luciferase activity was measured. (F) LLC-PK1 cells were transfected with the 1500PromBasic plasmid and SMAD2/SMAD3-siRNAs, after 24h the cells were treated with 20 mg/ml of albumin for another 24h. Luciferase activity was determined using a Luciferase Reporter Assay System, and the ß-galactosidase signal was used to normalize the results. Data are expressed as the means ± SD. Significant differences with controls are indicated by: *P ≤ 0.05; **P ≤ 0.01; ***P ≤ 0.001. Significant differences between treatments are indicated by # P≤0.05, ## P ≤ 0.01.

### The repression of megalin mRNA expression by TGF-ß1 is reverted by inhibition of histone deacetylases

Transcriptional repression involves, among other mechanisms, histone deacetylase (HDAC) activity [76]. We wanted to evaluate the effects of the HDAC inhibitor TSA on megalin mRNA expression in LLC-PK1 cells by qPCR. TSA treatment, starting at a 50 nM concentration, elicited a significant increase in megalin mRNA levels compared with the vehicle (control) (Fig 11A and 11B, third bar). These results are concordant with previous studies that analysed the regulation of megalin expression by TSA in NRK and Caco-2 cells [77]. We next examined the effect of TSA on TGF-ß1 -induced reduction of megalin expression that had been previously observed in LLC-PK1 cells (Figs 1A, 1B and 2A, 2B). The cells were treated with TGF-ß1 and/or TSA for 6 h and mRNA levels were then evaluated by qPCR. Whereas TGF-ß1 decreased mRNA as expected, TSA significantly counteracted TGF-ß1 response resulting in megalin mRNA levels similar to the control (Fig 11B). Interestingly, TSA alone also increases PAI-I expression, however, the induction of the PAI-1 gene by TGF-ß1 was blocked by TSA and was not additive, as has been described in other cell types [78, 79] (Fig 11C). In contrast to megalin mRNA levels, even though TSA alone increases megalin protein levels after 48 h of treatment, its effect over the reduction induced by TGF-ß1 was not evident under this experimental condition (Fig 11D). As a control for TSA treatment, the acetylated form of histone H3 was detected by immunoblotting after a 6 h treatment (Fig 11E). These results indicate that TSA counteracts the repressive effect of TGF-ß1 on megalin mRNA expression, via modulation of HDAC activity. However long term TGF-ß1 treatment may reduce megalin protein levels involving other pathways in addition or independently of the HDACs.

**Fig 11.**
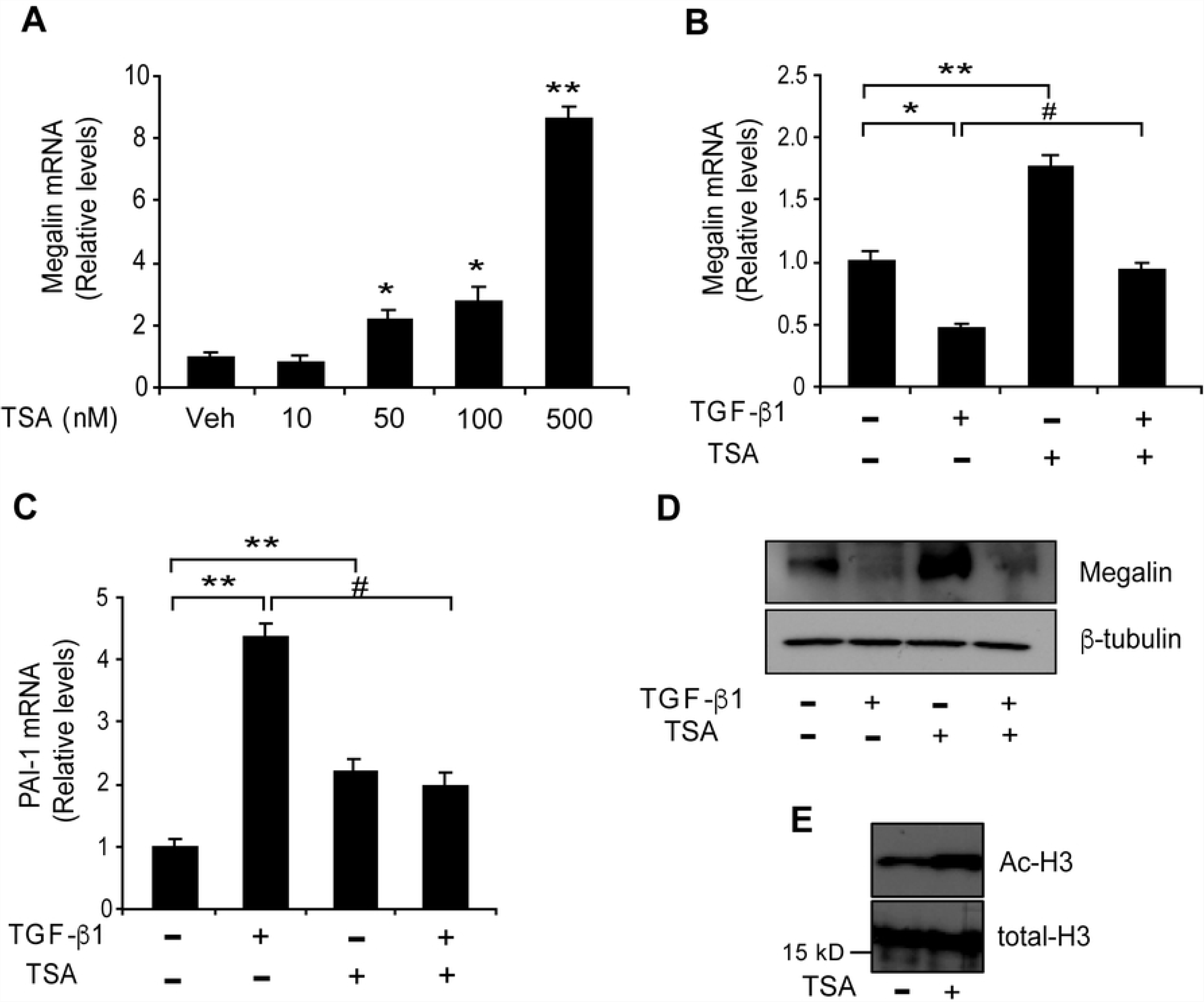
TSA increases megalin expression and counteracts the effect of TGF-β1 on megalin mRNA levels. (A) LLC-PK1 cells were treated with the indicated concentrations of TSA for 6 h, and megalin mRNA levels were analysed by real-time PCR. The cells were then exposed to TGF-ß1 (2.5 ng/ml) in the presence or absence of TSA (50 nM) for 6 h. Megalin (B) and PAI-1 (C) mRNA levels were then measured by real-time PCR. (D) LLC-PK1 cell extracts treated with TGF-ß1 (2.5 ng/ml) for 48 h in the presence or absence of TSA (50 nM) were used to measure megalin protein levels by immunoblotting; ß-tubulin was used as the loading control. (E) Cells were treated with TSA for 6 h. Total histones were extracted, and acetylated H3 and total H3 histones were detected by immunoblot analysis. The results are expressed as the means ± SD. Significant differences with the control (vehicle) are indicated by *P≤0.01, **P≤0.001. Significant differences with TGF-ß1 alone are indicated by #P≤0.001.

## Discussion

Due to the important roles of megalin already described in several systems in adult organisms and throughout development [27], understanding how the expression of this receptor is regulated has broad physiological significance. Among the factors that positively control megalin mRNA levels are retinoic acid [34, 80, 81], vitamin D [81], alpha and gamma peroxisome proliferator-activated receptor (PPAR) ligands [82] and bile acids [27, 35]. However, as it has already been discussed, megalin levels are reduced under fibrosis-associated conditions, i.e. diabetic nephropathy and gallstone disease [27, 47-49], ageing [83, 84] and cancer [34, 50], by mechanisms that are likely to involve TGF-ß as well as other molecules/mechanisms, including Ang II, leptin and albumin. However, the molecular mechanisms and signalling pathways associated with several of these negative regulatory processes are poorly understood.

Here, we show that the canonical SMAD-dependent TGF-ß1 signalling pathway directly regulates megalin expression. Although several lines of evidence link fibrotic processes with the direct or indirect effects of TGF-ß1 on megalin protein expression [47, 52], the molecular mechanisms underlying this reduction of megalin levels have not been determined. For example, in OK cells, interference with cellular SMAD2/3 activity by introducing dominant-negative SMAD2/3 resulted in partial abrogation of the TGF-ß1 - induced reduction of albumin internalization; the authors correlated this recovery in albumin uptake with the presence of megalin [52]. However, notably, albumin internalization can also be mediated by the neonatal Fc receptor [85]. Our study shows that the TGF-ß1 canonical signalling pathway directly represses megalin transcription, resulting in reduced megalin protein levels in two epithelial cell lines, renal proximal tubule LLC-PK1 cells and gallbladder dGBECs. By inhibiting the TGF-ßRI receptor (ALK5) [70], with SB-431542, we demonstrated that the observed decrease in megalin levels involves type I receptor activation. Interestingly, under our experimental conditions, megalin levels significantly increased when we used the inhibitor alone, suggesting that some of the TGF-ß1 -induced signalling pathways are active in this cell type and are responsible for the basal repression of megalin expression.

Using MatInspector software (www.genomatix.de) we analysed the human megalin promoter and found two SMAD protein response elements (SBEs) within the first 1,500 base pairs upstream of the transcription start site. Other SMAD-independent response elements, including Ap1, Sp1 and FAST1, are also present. Sp1 activity has been previously described for the human and rat megalin promoters [63, 86]. SMAD transcription factors form complexes that can simultaneously activate or repress hundreds of target genes in the same cell, depending on the context of the DNA-binding sites and on the presence of proteins that induce the formation of the protein complexes that bind to these sites [18]. In our study, immunoprecipitation assays revealed that SMAD2/3 bind to the megalin promoter at the two identified SBEs.

The functional role of these SBEs and of the SMADs in the repression of megalin expression induced by TGF-ß1, was deciphered using luciferase activation assays. The concomitant overexpression of transcription factors SMAD2 and SMAD3 caused a significant decrease in megalin promoter activity and also of megalin protein, in the same direction as the effect induced by TGF-ß1. These findings, combined with the results from promoter SBE mutations (double and single mutants) and the cellular knock-down of SMAD2/3, suggest that TGF-ß1 represses megalin promoter activation by regulating SMAD2/3 activity and binding to the megalin promoter SBEs.

Interestingly, the mutations in the SBEs led to a significant increase in relative luciferase activity, suggesting that the transcription factors SMAD2 and 3 basally suppressed megalin promoter activation. This result indicates that in LLC-PK1 cells SMAD2/3 are partially active. The observations of decreased megalin expression induced by the TGF-ßRI inhibitor SB-431542, and the presence of SMAD2/3 in the nucleus and the cytoplasm under control conditions that is reverted to the cytoplasm by SB-431542 treatment (Fig S1) support this idea. What could be the mechanism underlying this basal activation? Epithelial cells, including proximal tubule cells are able to secrete TGF-ß1 [51, 87, 88] and the effects of the cytokine relying on SMAD2/3 phosphorylation, could depend on the confluence level of the cells [89]. The presence of E-cadherin at cell-adhesions is associated with an inhibition of SMAD2/3 phosphorylation (and reduction of the signaling induced by TGF-ß1). In contrast, the reduction of E-cadherin localization at the membrane increases TGF-ß1 mediated SMAD phosphorylation [90]. In general, under the conditions in which we performed the activation experiments (mRNA expression, promoter activation and measurements of luciferase activity) the cells were not completely confluent as when we did the immunofluorescence studies (Fig 2B and 2D). If cells are not “superconfluent”, E-cadherin localizes both at the cell membrane and intracellularly. One possibility is that in our system these two elements (autocrine cytokine secretion and a partial E-cadherin mislocalization) are operating under basal conditions and therefore, we cannot discard that SB-421542 treatment is uncovering these mechanisms.

SMAD2/3 have the ability to induce gene activation because they interact with transcriptional co-activators such as p300 and CBP, which have histone acetyltransferase (HAT) activity [91]. Moreover, the SMAD proteins can also interact with transcriptional co-repressors such as TGIF [20], c-SKI [92], SIP-1 [93], SRF [94] and NF-1[95] to form complexes that can bind to DNA sequences such as FAST-2, TGF-ß inhibitory element (TIE), and SBEs, and can induce gene repression via HDAC recruitment. Regarding the possibility that these proteins might recruit HDACs, our work also shows that the repressive effect triggered by TGF-ß1 on megalin mRNA levels was counteracted by inhibition of HDAC activity. The inhibition of HDACs in non-stimulated cells increased megalin mRNA and protein expression; whereas long term TGF-ß1 treatment reduced megalin protein levels involving pathways not associated with HDACs. We cannot discard the possibility that other repressive mechanisms could be operating in the regulation of megalin by TGF-ß1. The human megalin promoter contains a CpG island that when methylated downregulates megalin expression [96]. Additionally, the inhibition of DNA methylation by 5Azacytidine increases megalin expression [77]. Moreover, recent evidence shows that TGF-ß1 regulates ECM production and differentiation controlling gene expression through DNA and histone methylation [97]. TGF-ß1 also regulates EMT in cancer cells through an increase in DNA methylation and histone 3 lysine methylation at the E-cadherin promoter, depending on DNA methyltransferase 3A [98].

Another system that decreases megalin expression is induced by exposure to a high concentration (10-20 mg/ml) of albumin [57], as we showed here and as has been shown previously [82]. Although the mechanisms involved in the reduction of megalin expression by albumin are not entirely clear, other studies have demonstrated that albumin can induce the release of TGF-ß1 in PTCs [99]. Furthermore, albumin can directly activate the TGF-β1 type II receptor in endothelial cells [100] and TGF-β1 secretion in astrocytes [101] via SMAD2/3 pathway activation [102]. Moreover, albumin has been shown to induce TGF-ß1 expression through the EGF receptor and ERK-MAPK signalling activation after a 72 h treatment [74]. In the present study, we found that albumin reduces megalin mRNA expression and activation of its promoter. The inhibition of TGF-ß1R activation as well the decrease in SMAD2/3 cellular levels significantly counteracted the effect of albumin on megalin expression, reinforcing the idea that the SMAD2/3-TGF-ßRI canonical signalling pathway mediates part of the effect of albumin on megalin levels.

In addition to albumin, other megalin ligands such as Ang II, leptin and CTGF induce fibrotic responses through their signalling receptors [55, 103-106]. Ang II, through its signalling receptor, AT1, may also stimulate the SMAD-dependent pathway during the EMT, causing renal fibrosis [103, 106]; and leptin also induces TGF-ß1 expression [51]. As is the case for high albumin concentrations, both Ang II and leptin have been associated with reduced megalin expression [51, 58]. For leptin, the reduction in megalin was not observed at the transcriptional level; thus, the authors suggested that this reduction was an acute effect that could likely be explained by megalin degradation. However, considering our results, chronic effects could include the regulation of megalin transcription because the expression of TGF-β1 is increased by leptin.

In this study, we showed that TGF-ß1 at low concentrations (2.5 to 10 ng/ml) reduced total megalin protein levels only as a result of decreased transcription and not of increased receptor degradation. Megalin half-life showed no changes in cells treated with TGF-ß1 at two different concentrations (2.5 and 20 ng/ml) and in the window of time considered in our study (up to 12 h). Importantly, the concentrations of TGF-ß1 used in most of our work (2.5 to 10 ng/ml) are in the physiological range but the concentration of 20 ng/ml is rather high and in the range of diabetic patients [107, 108] In addition, the cytokine did not modify receptor levels at the cell surface as could be expected by the shedding of the ectodomain. Therefore, TGF-ß signalling did not increase receptor degradation within the first 12 h of treatment. Recently, Mazzocchi et al [73] showed that in alveolar cells, TGF-ß1 is able to reduce surface megalin levels by inducing shedding of the megalin ectodomain, specifically by TGF-ß-induced secretion of matrix metalloproteases (MMPs)−2, −9 and −14. This process is followed by γ-secretase regulated intramembrane proteolysis (RIP) that induces the generation of the MICD as described previously [59, 109]. The authors indicate that only high TGF-ß1 concentrations (20 ng/ml) decreased megalin mRNA levels after 24 and 48 h treatments, suggesting that this effect was the result of the auto inhibitory effect of MICD as repressor of megalin transcription. In contrast with the report of Mazzocchi et al, our study did not find an increase in the levels of megalin CTF, the MICD precursor, after 10 h of TGF-ß1 treatment. Moreover, we found a significant decrease in megalin mRNA expression already detectable after 5 h and with a cytokine concentration almost 10 times lower than the one used on the alveolar cells. We cannot discard that, in addition to our observation that TGF-ß1 directly represses megalin transcription via SMAD2/3, the MICD-dependent mechanism could also be operating. However, this mechanism could only function with a 10-fold increase in the concentration of the cytokine and in a different time frame. TGF-ß1 also induces the proteolytic shedding of E-cadherin [4], an epithelial membrane protein that was significantly decreased by TGF-ß1 in our study. The E-cadherin reduction induced by TGF-ß1 could also be caused by gene repression mediated by SMAD-interacting protein (SIP-1) via co-repressor C-terminal binding protein (CtBP) recruitment [110]; as well as by the activation of different transcription factors such as Snail, Twist, Slug, and Zeb that bind to specific sequences (called E-boxes) within the promoter [111].Therefore, more than one mechanism could be activated in response to TGF-ß1 exposure that results in functional reduction of megalin.

Considering the role of megalin in regulating the availability of fibrotic molecules, the decrease of megalin expression induced by its own ligands (and by TGF-ß) could be associated with the evolution and/or the severity of some pathological conditions, such as fibrosis and cancer, in different megalin-expressing epithelial tissues. In other words, megalin itself could be considered a protective agent for those conditions by acting, for example, as an anti-fibrotic protein. Similarly, in some cancer cells, megalin expression is also decreased [34, 50]. Because megalin is required for the internalization and activation of Vitamin D to 1,25-OH Vitamin D [45, 46], its decrease would directly impact the activation of its nuclear receptor, VDR, which plays an important anti-proliferative role in the control of some cancer types, especially breast, prostate and colon cancer [112].

## CONCLUSION

The knowledge generated in this work establishes the molecular mechanism involved in the reduction of megalin expression by TGF-ß, and will contribute to understanding in more detail the regulation of megalin in diseases in which this pro-inflammatory cytokine plays a central role.

## ACKNOWLEDGEMENTS

The authors thank Dr. Rifkin (NYU School of Medicine, NY, USA) for providing the p800Luc plasmid. We also thank Dr. J. F. Miquel and Lorena Azocar (Faculty of Medicine, Department of Gastroenterology, Pontificia Universidad Católica de Chile) for their technical support throughout this work. We thank Dr. Enrique Brandan (Faculty of Biological Science, Department of Cellular and Molecular Biology, Pontificia Universidad Católica de Chile) for providing us with the SB-431542 inhibitor, Dr. Miguel Bronfman (Faculty of Biological Science, Department of Cellular and Molecular Biology, Pontificia Universidad Católica de Chile) for providing some equipment, Dr. Frank Lammert (Department of Internal Medicine I, University Hospital Bonn, University of Bonn, Bonn, German) for the −1500 pGL3 basic megalin promoter plasmid (−1500PromBasic), Dr. Sum P. Lee (Department of Medicine, University of Washington, and VA Puget Sound Health Care System, Seattle, WA) for the generous gift of the canine GBEC line, Drs. María Estela Andrés and Montserrat Olivares (Faculty of Biological Science, Department of Cellular and Molecular Biology, Pontificia Universidad Católica de Chile) for their invaluable technical contributions for the ChIP assays and Gabriela Valarezo for her contribution in some biotinylation experiments and manuscript editing.

## Author contribution statement

**FC** and **PF** performed most of the experiments and designed some of the figures. **MPM** conducted the research and wrote the manuscript.

